# Simultaneous Identification of Changepoints and Model Parameters in Switching Dynamical Systems

**DOI:** 10.1101/2024.01.30.577909

**Authors:** Xiaoming Fu, Kai Fan, Heinrich Zozmann, Lennart Schüler, Justin M. Calabrese

## Abstract

Many complex natural systems undergo shifts in dynamics at particular points in time. Examples include phase transitions in gene expression during the cell cycle, introduced species affecting predator-prey interactions, and disease outbreaks responding to intervention measures. Such changepoints partition timeseries into different dynamical regimes characterized by distinct parameter sets, and inference on both the changepoints and regime-specific dynamical parameters is of primary interest. Conventional approaches to analyzing switching dynamical systems first estimate changepoints, and then estimate dynamical parameters assuming the changepoints are fixed and known. Such two-stage approaches are ad-hoc, can introduce biases in the analysis, and do not fully account for uncertainty. Here, we introduce a rigorous, simulation-based inference framework that simultaneously estimates changepoints and model parameters from noisy data while admitting full uncertainty. We use simulation studies of oscillatory predator-prey dynamics and stochastic gene expression to demonstrate that our method yields accurate estimates of changepoints and model parameters together with appropriate uncertainty bounds. We then apply our approach to a real-world case study of COVID-19 intervention effects, and show that our inferred changepoints aligned closely with the actual dates of intervention implementation. Taken together, these results suggest that our framework will have broad utility in diverse scientific domains.

## Introduction

Real-world complex systems can experience dramatic shifts in their dynamical behavior due to internal feedbacks or changes in external conditions. For example, gene expression undergoes phase transition during the cell cycle driven by cell size growth and cellular signaling pathways [1, 2, 3, 4], and physical and biological systems experience phase transitions under non-equilibrium dynamics [5]. Importantly, the effects of human actions, either inadvertent or intentional, can also lead to shifts in system dynamics at particular points in time. One classic example is lake eutrophication, whereby increased inputs of phosphorous from fertilizers drive a transition from a clear water oligotrophic state to a turbid eutrophic state [6]. A more recent example is COVID-19 outbreak dynamics responding to governmental non-pharmaceutical interventions (NPIs) [7, 8]. Indeed, examples of switching dynamical systems arise across the sciences, and in both basic research and applied contexts.

Switching dynamical systems can be modeled via (possibly coupled) differential equations with state variables and/or model parameters that change values at particular times [9]. By definition, the system’s behavior shifts substantially at such changepoints, which affects the overall dynamics of the system. Previous work on such state variable discontinuities have been successfully applied to a variety of settings. Examples include ordinary/stochastic differential equations with explicit events [10, 11] and neural ordinary differential equations with implicit events [12]. However, to the best of our knowledge, a simultaneous inference framework for the changepoints and parameter discontinuities in switching dynamical systems has not been developed.

When analyzing timeseries from switching dynamical systems, shifts in system behavior can be modeled statistically by segmenting timeseries at identified changepoints into sections characterized by different parameter sets. Typically, both the changepoints themselves as well as the segment-specific dynamical parameters are targets of inference. Even when only a subset of the parameters is of interest, all parameter estimates must be appropriately (and simultaneously) estimated to avoid both biases and underestimation of uncertainty. Unfortunately, conventional parameter estimation approaches for dynamical systems with changepoints typically proceed in two steps [13, 14, 15, 16]. First, changepoints are estimated using statistical methods based on (generalized) linear models or Bayesian regression [17, 18, 19, 20]. Second, the changepoints thus identified are treated as fixed and know, and then dynamical parameters are estimated separately for each resulting segment of the timeseries. While conceptually straightforward, such two-stage approaches are ad hoc and fail to account for uncertainty in estimated changepoints when quantifying uncertainty in dynamical parameters, which can lead to overly-confident inference on parameters of interest. Furthermore, treating changepoints as fixed can also lead to biased estimates of dynamical parameters.

Here, we introduce a simulation-based framework for simultaneous estimation of changepoints and model parameters in nonlinear switching dynamical systems. Our approach is applicable to a large group of models governed by reaction rate laws derived from underlying stochastic processes, e.g. ordinary differential equations (ODEs) based on mass-action kinetics [21, 22]. In addition, our approach accommodates scenarios where the analytical likelihood function is mathematically intractable by incorporating a likelihood approximation component. All these points provide a flexible and computationally efficient approach, fully accounts for uncertainty in both changepoints and model parameters, and can be used to support either likelihoodist or Bayesian inference.

Taking a Bayesian approach, we demonstrate the superior performance of our methods on three contrasting case studies. First, we investigate a predator-prey system (Section Case Study: Lotka-Volterra (Predator-Prey) Model), in which the intrinsic traits of periodic oscillatory solutions are often challenging for conventional two-stage approaches. This example serves to demonstrate the inferential accuracy and robustness of our method over two-stage approaches when applied to a highly nonlinear system. We then apply our method to an asynchronous stochastic gene expression model (Section Case Study: Stochastic Gene Expression Model). This model represents an ideal case where two-stage methods should perform well because this model admits a closed-form likelihood function and, in our simulations, features long runs of distinctly different dynamics on either side of a single change point. We select this example to evaluate the performance of our proposed method under conditions favorable to the two-stage approach. Finally, we study an epidemiological model (Section Case Study: Compartmental Models in Epidemiology) to demonstrate the utility of our method in real-world applications involving limited, noisy data. Notably, when applying our method to analyze COVID-19 data from Germany, the estimated non-pharmaceutical intervention (NPI) changepoints are closely aligned with the actual NPI dates. In all cases, our method outperforms conventional two-stage approaches to estimating the parameters of switching dynamical systems, and thus represents a significant advance in the analysis of this diverse, important, and widely used class of complex systems.

## Results

Suppose we are interested in estimating the changepoints and model parameters of an ODE system

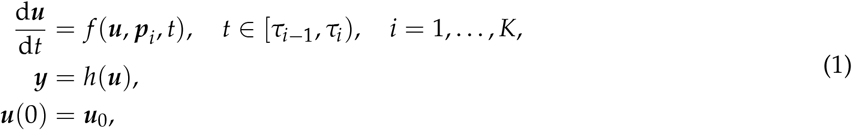

where the parameter of the system changes from ***p***_*i*_ to ***p***_*i*+1_ at the change points *τ*_*i*_ for *i* = 1, …, *K* 1 (set *τ*_0_ = 0, *τ*_*K*_ = *t*_end_). Here ***y*** represents observable variables of the system, which can be obtained by applying a mapping *h* to the state variables ***u***. Given a set of observed data 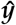 and an objective function 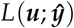, the goal is to find the optimal model parameters 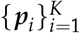 and changepoints 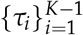 such that the objective function is minimized. Hence, any gradient-based optimization method requires the gradient of the objective function with respect to the model parameters and changepoints.

Inspired by the work of [23, 24, 10], we derive the adjoint equations to compute the gradients for the changepoints and model parameters. This technique, known as reverse-mode sensitivity analysis, calculates how changes in parameters affect the final output of a system by solving the adjoint equations backward-intime (similar to the backpropagation in training a neural network). Suppose the objective function is given by 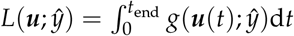 d*t* where *g* is a general scalar-valued function and ***u***(*t*) is the solution of the ODE system (1). The sensitivity of the changepoint and model parameters (see Methods section Differentiating Through Changepoints and Model Parameters for details) can be calculated by:

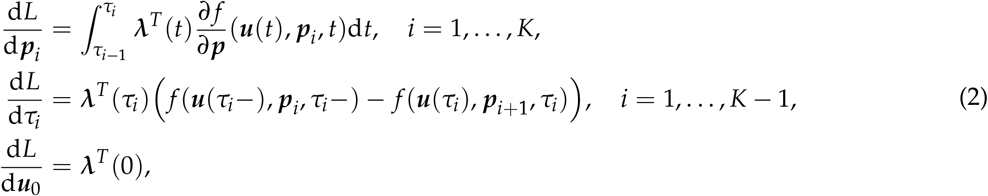

where *τ*_*i*_− denotes the left limit and ***λ***(*t*) satisfies the adjoint equations

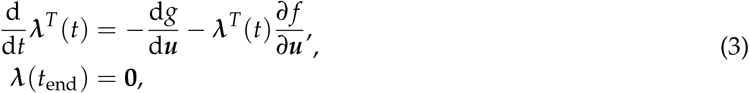

solving backward from *t*_end_ to 0.

The sensitivity analysis serves for obtaining the gradient information of the changepoint and parameters which can be used for both maximum likelihood estimation (MLE) and Bayesian inference using Hamiltonian Monte Carlo (HMC) algorithms (see Methods section Bayesian Inference with Hamiltonian Monte Carlo for more details). A Bayesian approach for parameter estimation requires prior knowledge about reasonable parameter ranges and the likelihood function. The posterior distribution, obtained by updating our prior knowledge about model parameters with data, is generally narrower than the prior. This is the well-known formula for the posterior according to Bayes’ rule

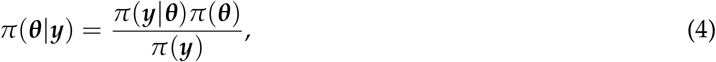

where *π*(***y*** |***θ***) quantifies the likelihood of observing data ***y*** given the parameters ***θ***, *π*(***θ***) is the prior distribution capturing our knowledge about plausible parameter configurations for ***θ***, and the denominator represents a normalizing constant (usually called the evidence). Using the HMC sampling method we can update the posterior *π*(***θ*** |***y***) without explicitly calculating the evidence *π*(***y***), thus the key step for the inference is to calculate the likelihood *π*(***y***| ***θ***).

In fact, if ODE system (1) is based on the law of mass action [25], then its meso-scopic representation (known as stochastic reaction networks [SRNs]) can be used to derive or approximate the likelihood (see Methods section Constructing the Likelihood Using the Linear Noise Approximation). Some early works considered approximating the likelihood by using transition probabilities of the continuous-time Markov process of the SRNs from the Langevin equation [26]. Other works proposed inference schemes which utilize the linear noise approximation (LNA) [27, 28]. LNA provides sufficient statistics of the (Gaussian) single-time marginal distributions, and can be efficiently computed by integrating a system of ordinary differential equations [29]. In this work, we leverage the efficient computation of the LNA to calculate the likelihood, which can then be used for Bayesian inference.

### Case Study: Lotka-Volterra (Predator-Prey) Model

The Lotka-Volterra model describes the dynamics of predator-prey interactions in an ecosystem. It assumes that the prey population *U* grows at a rate represented by the parameter *α* in the absence of predators, but decreases as they are consumed by the predator population *V* at a rate determined by the interaction strength parameter, *β*. The predator population decreases in size if they cannot find enough prey to consume, which is represented by the mortality rate parameter, *δ*. The corresponding ODE system reads

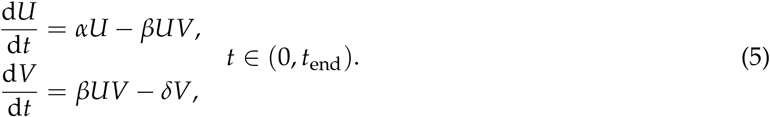

When a new prey species is introduced, it can compete with the existing prey for resources, leading to a decline in the growth rate *α* of the original species. This competition can also weaken the interaction strength *β* between the predator and prey populations, as predators can now switch to the new prey. Such a change can be modeled by introducing a changepoint *τ* into the system. Hence, both the growth rate of prey population *α* and the interaction rate *β* are assumed to be piecewise constant functions, that is

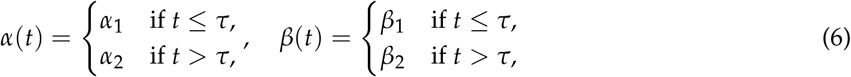

where *τ* is the changepoint and 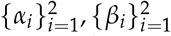 are the corresponding piecewise rates. While the mortality rate *δ* is also a parameter to be estimated, it is assumed to be constant over the entire timeseries.

Such a system can be modelled by a SRN to study the probability of the system being in a particular state at a given time described by the master equations (MEs) [30] (see Methods section Constructing the Likelihood Using the Linear Noise Approximation for details). While the Lotka-Volterra model lacks an analytical solution to the corresponding MEs, we can use the LNA to obtain the approximate mean and variance of the Markov process generated by the SRN. Hence, the likelihood function of the predator-prey model is regarded as a normal distribution with time-dependent mean and variance given by the solution of the LNA system (see Supplementary Information A for details).

For the parameter inference, we draw *N* sets of random model parameters 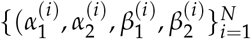 and change points 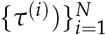 from a set of sample distributions with *N* = 300. Each synthetic timeseries data is generated based the ODE system using one parameter set (see Supplementary Information A). The inferential task is to accurately estimate both the change points and model parameters from the synthetic data. The results are shown in Figure 1 **c** and **d**.

**Figure 1:**
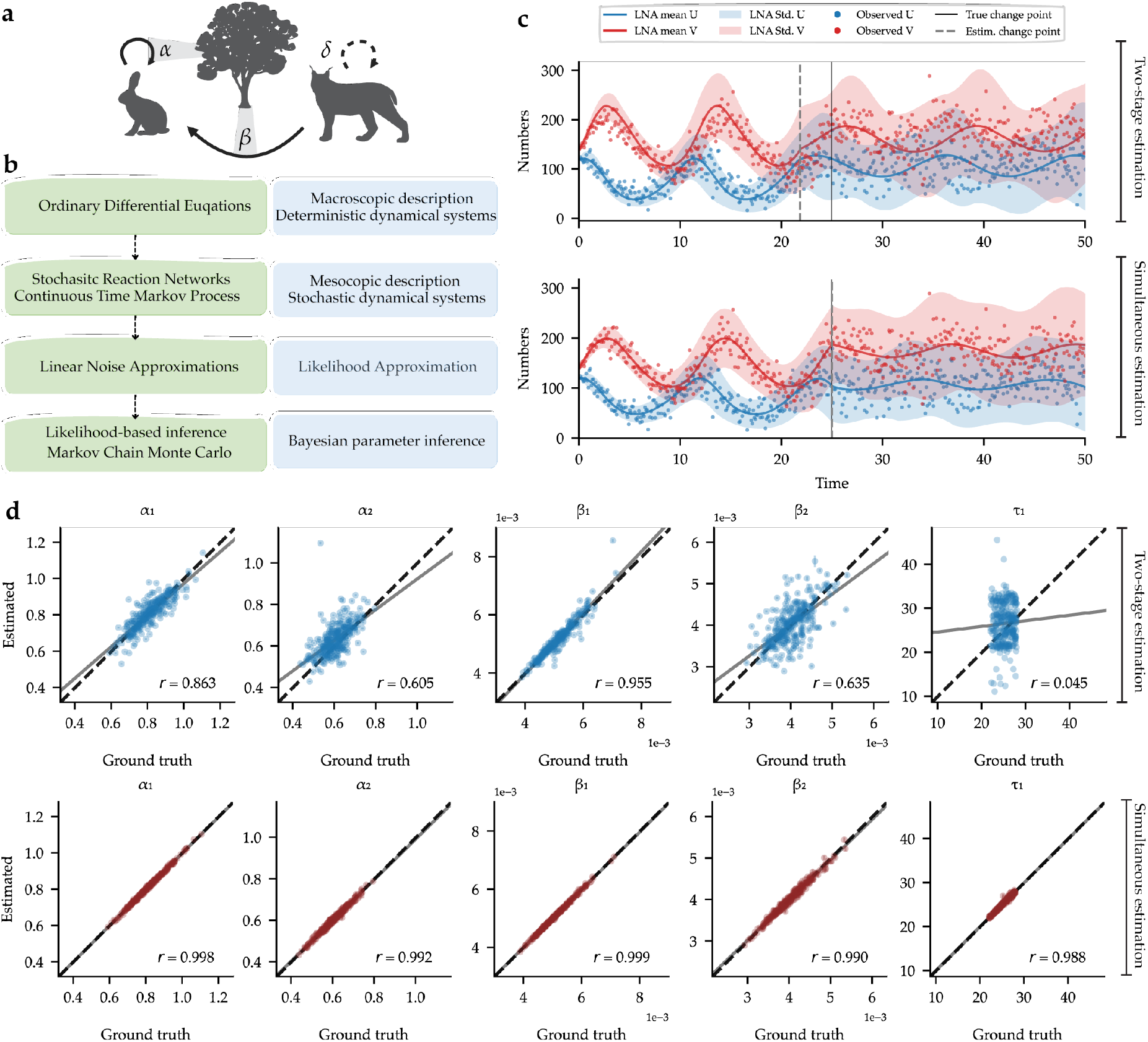
Bayesian inference based on the LNA applied to the predator-prey system. **a**. Sketch of a predator-prey system: the prey population grows at a rate represented by *α* but decreases as they are consumed by the predator population with rate *β*. These two parameters undergo a change point due to environmental changes, while we assume the mortality rate *δ* of the predator population is not affected by the change. **b**. The inference framework: starting from the macroscopic description of the population dynamics, we consider its stochastic representation (SRN) and use the LNA to estimate the population’s mean and (co)variance for the likelihood function in the Bayesian inference. **c**. Comparing mean and variance of the LNA to the predator-prey system with estimated parameters using an ad-hoc two-stage method and our simultaneous estimation method, respectively. The synthetic data is generated based the ODE system, and shaded areas represent the standard deviation of the LNA solution. The ground truth and estimated change points are shown in solid and dashed lines respectively. **d**. Inferential accuracy computed from 300 simulated predator-prey scenarios using the best-performing two-stage method and our simultaneous method. Perfectly accurate estimates fall on the 1:1 line (black, dashed), while the ordinary least square linear regression lines (gray, solid) show the trend observed in the estimates. The Pearson product-moment correlation coefficients (*r*) are shown in each subfigure.

Specifically, in Figure 1 **c**, our focus narrows down to examine one representative set of inference results among the 300 realizations. We compare the solutions generated by the median of the posterior samples using both the best-performing two-stage method and our simultaneous estimation method. For both methods, the standard deviation of the LNA solution adequately covers the observed data. The precision of the changepoint estimate achieved by our method surpasses that of the two-stage method when comparing to the true changepoint. Consequently, the mean of the LNA solution, calculated using our proposed methodology, demonstrates a greater fidelity to the ground truth in contrast to the two-stage method (with negative log-likelihood: 4787.9 (two-step) vs 4530.2 (ours)).

Figure 1 **d** compares the estimated median of posterior samples to the ground truth, with the error bars marking the median absolute deviation of the posterior samples. The scatter plot shows that the majority of the data points are situated close to the 1:1 lines using our method, indicating a strong correlation between the estimates and associated ground truth values. Furthermore, the high Pearson product-moment correlation coefficients (*r* values), both exceeding 0.9, demonstrate an accurate estimation of the parameters. However, the two-stage method shows a large deviation from the 1:1 lines, indicating a poor correlation between the variables with significantly lower *r* values.

Furthermore, to check the ability of our Bayesian model to recover the correct target posterior, also referred as computational faithfulness, we use simulation-based calibration (SBC) [31]. SBC re-uses the 300 simulated predator-prey scenarios. The uniformity of the rank statistics indicates that there are no systematic biases in the approximate posteriors and implies trustworthy approximation (see Supplementary Information Figure 4 for details).

Lastly, further comparisons with the two-stage inference method over synthetic data are presented in Supplementary Information D. The results show that two-stage methods are generally not able to capture the uncertainty in the model parameters, and thus are not able to recover the correct target posteriors. A comparison with a likelihood-free method using approximate Bayesian computation (ABC) is also presented in Supplementary Information A.

### Case Study: Stochastic Gene Expression Model

Next, we consider a birth-death stochastic gene expression model with changepoints. The stochastic reaction network is given by

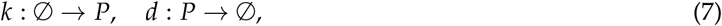

where *k* : ∅ → *P* represents that the product *P* (e.g. protein) is produced at a time-dependent initiation rate *k*, with ∅ representing the empty reactant because how *P* is produced is not of interest here. Similarly, *d* : *P* → ∅ represents that *P* leaves the system of interest at degradation rate *d*. The corresponding MEs are given by:

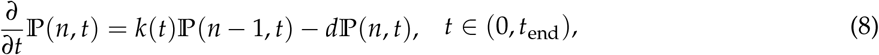

where ℙ(*n, t*) is the probability that the system is in state *n* at time *t* (see Methods section Constructing the Likelihood Using the Linear Noise Approximation for derivation). The production rate *k* is assumed to change at the changepoints *τ*, that is

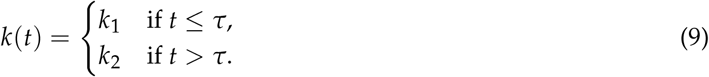

It is well-known that during a cell cycle the dynamics of stochastic gene expression can undergo some phase transition, thus affecting the effective initiation rate *k* [4, 32, 33]. More importantly, due to cell-to-cell variability, such a transition is asynchronous in a population, that is, the transition happens at different times with different initiation rates for different cells [34, 35]. In this example, we introduce the heterogeneity of the cell dynamics by assuming that the initiation rates *k*_1_, *k*_2_ and the changepoint *τ* are random variables for each cell (see Figure 2 **a** for illustration).

**Figure 2:**
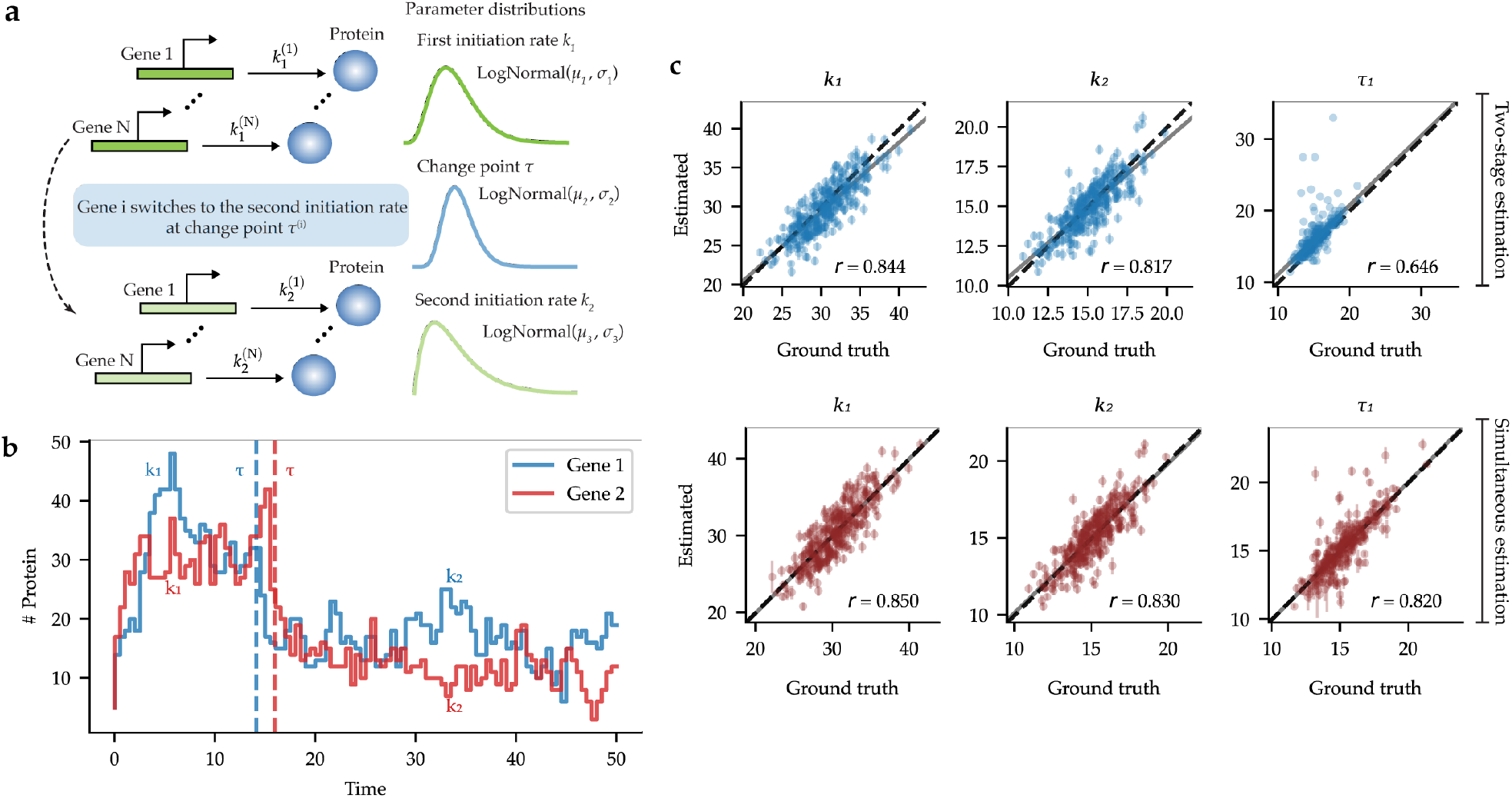
Recovering the underlying distributions of the kinetic parameters of a stochastic gene expression model. **a**. Sketch of a stochastic gene expression model: each gene *i* initially produces a protein at a fixed initiation rate 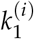 and degrades at rate *d* until change point *τ*^(*i*)^, after which the initiation rate switches to 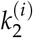. Both the model parameters 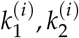 and the change point *τ*^(*i*)^ are drawn from log-normal distributions to mimic the heterogeneous and asynchronous features in a cell population. **b**. Two random trajectories with the asynchronous change points and different model parameters. **c**. Comparison of the ground truth and estimated parameters under Bayesian inference using two-stage inference method and our inference method. The Pearson product-moment correlation coefficients (*r*) are shown in each subfigure and dashed (black) and solid (gray) lines are as in Fig. 1.

For this simple example with one changepoint, we can derive that the single-time solution of the MEs (8) follows a Poisson distribution, that is,

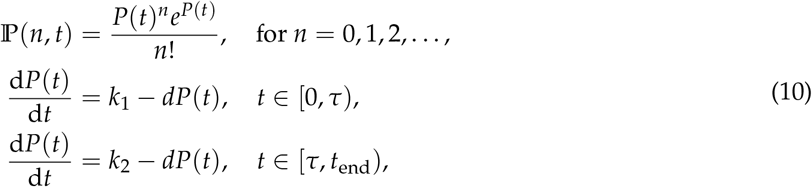

where *P*(*t*) is the solution of the corresponding ODE system. Hence, the analytical solution to the MEs (10) is used as the likelihood function for Bayesian inference.

For inference on model parameters, we draw 300 sets of random parameters 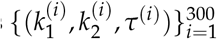 from the prior distributions. For each parameter set, we use the stochastic simulation algorithm (SSA) [36] to simulate a random trajectory from the SRN of the gene expression model (7) (see Supplementary Information B for details). The resulting parameter estimates are presented in Figure 2.

As we can see in Figure 2 **b**, the distinct change points evident in the trajectories render them amenable to analysis using the conventional two-stage method. Each segment of the trajectory exhibits the characteristics of a simple birth-death process, and there are sufficient data points on either side of the changepoint to facilitate reliable estimation of kinetic parameters. Indeed, we selected this example to evaluate the performance of our proposed method under conditions highly favorable to the two-stage approach.

As shown in Figure 2 **c**, the results of both methods show a strong correlation between the inferred and ground truth model parameters *k*_1_, *k*_2_. The Pearson correlation coefficients (*r*) are above 0.8, indicating a reliable estimation. However, the two-stage method exhibits a substantial deviation for the changepoint estimation *τ*, with a Pearson correlation coefficient of 0.646. In contrast, our method demonstrates superior accuracy with a Pearson correlation coefficient of 0.820. This enhanced accuracy stems from our method’s ability to effectively capture all the uncertainty in the model parameters and changepoints, which is not possible with the two-stage method (see Supplementary Information D for more details). More details on the numerical implementations can be found in Supplementary Information B. Thus even in a case that plays to the strengths of two-stage approaches, our simultaneous estimation method still exhibits superior performance.

### Case Study: Compartmental Models in Epidemiology

After validation of the general applicability and accuracy of our inference method on simulated data, we now consider a real-world case study from epidemiology. In particular, we model the spread of COVID-19 in Germany during the early months of the pandemic based on data from the federal agency tasked with disease control and prevention in Germany. In epidemiology, compartmental models are widely used to describe the spread of infectious diseases. One of the simplest models is the SIR model, which consists of three compartments: susceptible, infected and recovered. Here we consider a SEIRD (susceptible, exposed, infected, recovered and deceased) model [37, 38] as an extension including exposed (but not infectious) and deceased compartments, which is given by

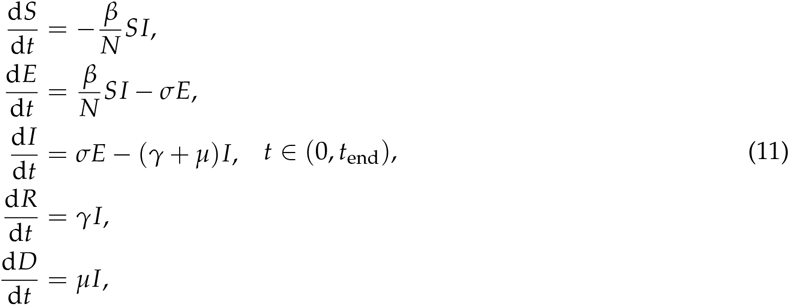

where *β* is the transmission rate, *γ* is the recovery rate and *N* is the total number of the population. The parameter *σ* is the rate of exposed individuals becoming infectious, and *μ* is the rate of infected individuals dying due to infection.

The model is applied to 120-day span of epidemiological data on the number of reported COVID-19 cases (infected and deceased) in Germany from March 1^st^, until June 28^th^, 2020. The data were obtained from publicly available sources [39] are not subsequently cleaned or corrected in the aftermath. This example was chosen specifically because of the knowledge of the changepoints (based on NPI implementation dates and appropriate reporting delays), providing an objective check on the performance of the different methods with respect to changepoint estimation.

The changepoint is defined as the time when the transmission rate, *β*, changes in response to the implementation of a particular non-pharmaceutical intervention (NPI). In reality, this may correspond to a variety of response measures introduced by the government to contain transmission of COVID-19, such as the closure of public spaces and stay-at-home orders. Additionally, we assume that the mortality rate *μ* is impacted by changes in hospital admission policies, such as the availability of hospital beds increasing over time and the reporting of more COVID-related deaths. Thus, we assume that the transition rate from the infected to the deceased compartments, *μ*, is a piecewise constant function, but the changepoints can be different from those of *β*, that is

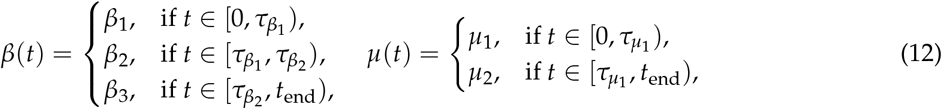

where 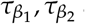 and 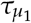 are the changepoints for *β* and *μ*, respectively. Note that here we set two changepoints for *β* and one for *μ*. In fact, our inference suggests that assuming two sets of changepoints for both *β* and *μ* gives similar results (see Supplementary Information C for details).

The transition rate from the exposed to infected class, *σ*, is also a parameter to be estimated but is assumed to be constant over time. In contrast, we fix the recovery time for the original variant to *γ* = 1/14 day^−1^ based on literature estimates between (12.6, 20.6) days (CI 95%) [40]. Figure 3 **a** shows the schematic diagram of the SEIRD model with changepoints. For the inferential accuracy check using synthetic data, the prior distributions and implementation details, please see the Supplementary Information C, as well as for a comparison study with two-stage estimation D.

**Figure 3:**
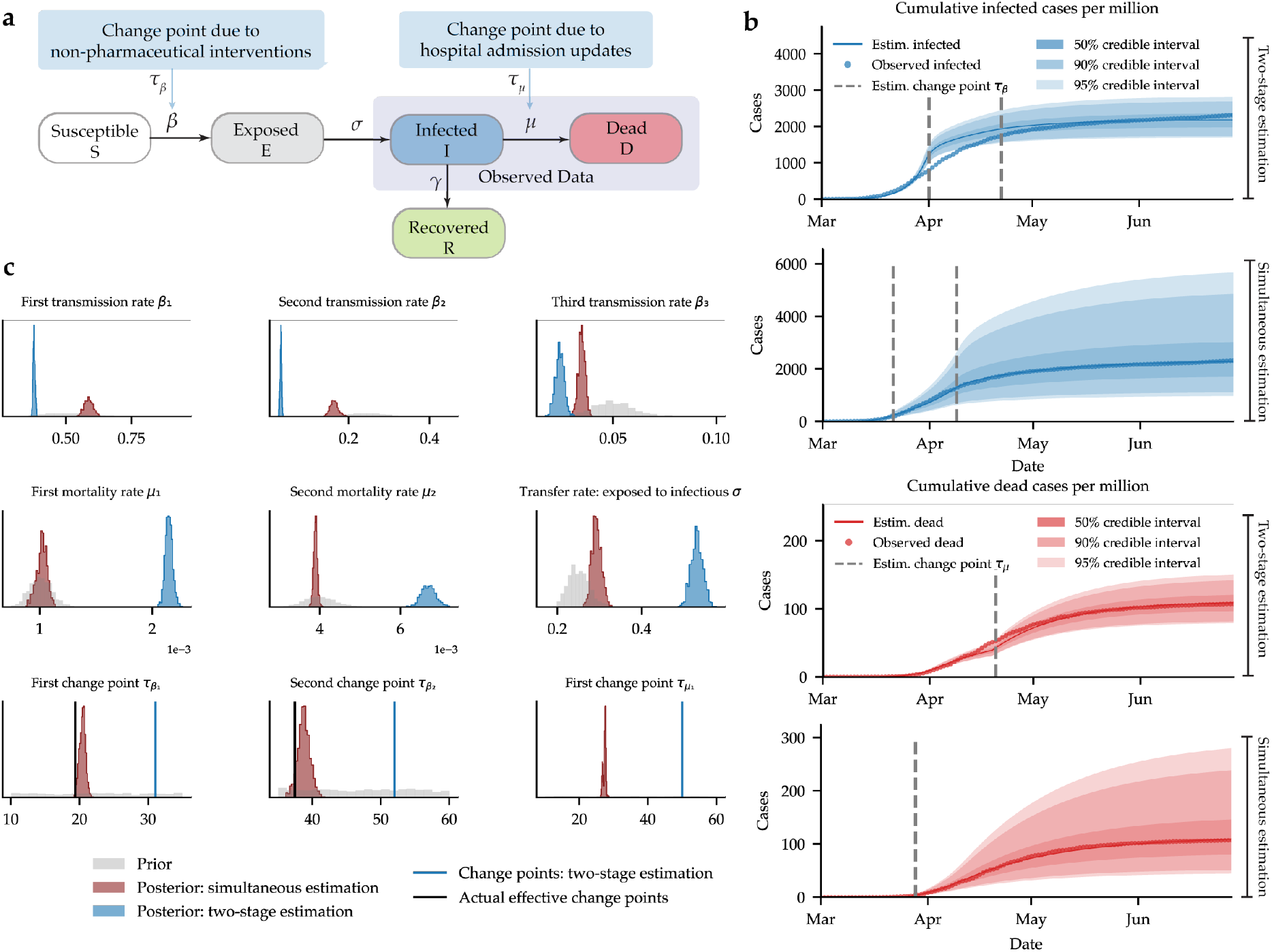
Inference results based on the SEIRD model for reported COVID-19 cases and deaths in Germany. **a**. The schematic diagram of the SEIRD model with one set of changepoints on the transmission rate *β* related to the NPI implementation, and a separate changepoint on the mortality rate *μ* related to the hospital admission updates. **b**. Posterior predictions of new cases obtained by inferring model parameters from epidemiological data available for reported cases and deaths under the best-performing two-stage method and our simultaneous estimation approach. The estimated changepoints are presented as dashed vertical lines. The shaded areas represent the credible intervals (CI) of the posterior predictions. **c**. Marginal posteriors of all model parameters inferred from data under the best-performing two-stage method and our simultaneous estimation method. Gray histograms depict prior distributions for comparison with the posteriors (red: our simultaneous methods; blue: the best-performing two-stage method). In the estimated changepoints, blue vertical lines indicate the estimated change points using two-stage method. Black vertical lines indicate the actual effective dates of NPIs in Germany. The full details of the estimated parameters are shown in Table 1.

Posterior predictions of the SEIRD model for the infected and dead in Germany are shown in Figure 3 **b**. Using our proposed method, standard point estimates (median) and credibility intervals (50%, 90% and 95% CI) are shown, with the reported cases always lying within the estimated CIs. The estimated change points are presented as dashed vertical lines. Conversely, while the two-stage method produces narrower posterior predictions, it fails to adequately capture the trend of the data. This discrepancy can be attributed to two causes: (1) the estimated changepoints deviate from the actual NPI dates; and (2) dynamical parameter estimates that are both biased and overly confident, resulting in inaccurate predictions of reported cases. This comparison underscores the superior performance of our proposed method in accurately inferring all model parameters, including changepoints, and in providing more reliable predictions. The full details of the estimated parameters are shown in Table 1.

**Table 1:** Parameter estimation results for the German COVID-19 data with changepoints in the transmission and mortality rates.

| Description | Parameter | Mean | Median | MAP | 95%-CI |
| --- | --- | --- | --- | --- | --- |
| Transmission rate $S \rightarrow E$ (1/days) | $\beta_1$ | 0.59 | 0.59 | 0.58 | [0.55, 0.62] |
| | $\beta_2$ | 0.17 | 0.16 | 0.16 | [0.14, 0.19] |
| | $\beta_3$ | 0.03 | 0.03 | 0.03 | [0.03, 0.04] |
| Mortality rate $I \rightarrow D$ ($1 \times 10^{-3}$/days) | $\mu_1$ | 1.01 | 1.02 | 1.03 | [0.92, 1.10] |
| | $\mu_2$ | 3.88 | 3.89 | 3.87 | [3.80, 3.98] |
| Incubation rate $E \rightarrow I$ (1/days) | $\sigma$ | 0.29 | 0.29 | 0.28 | [0.27, 0.32] |
| Change point for the transmission rate | $\tau_{\beta_1}$ | Mar-21<br>(Day 20.42) | Mar-21<br>(Day 20.53) | Mar-21<br>(Day 20.53) | [Mar-20, Mar-22]<br>[Day 19.52 - 21.30] |
| | $\tau_{\beta_2}$ | Apr-09<br>(Day 38.72) | Apr-09<br>(Day 38.67) | Apr-09<br>(Day 38.67) | [Apr-07, Apr-10]<br>[Day 36.87 - 40.46] |
| Change point for the mortality rate | $\tau_{\mu_1}$ | Mar-28<br>(Day 27.37) | Mar-28<br>(Day 27.46) | Mar-28<br>(Day 27.46) | [Mar-27, Mar-29]<br>[Day 26.44 - 27.82] |

Our analysis shows that despite the large number of unknown parameters and available data on only two compartments (namely, infected and deceased groups), there is a significant reduction in uncertainty in relation to the prior knowledge for all model parameters (Figure 3 **c**) for our simultaneous estimation approach. If we subtract the median of the implementation delay of around 14 days (estimated change duration plus the reporting delay according to [7]), our first estimated NPI was on March 6^th^ (CI [5, 7]), 2020. This matches the cancellation of large public events (trade fairs and soccer matches). Our second estimated NPI was on March 24^th^ (CI [22, 26]), 2020. This corresponds to the strict contact ban and further response measures, which were implemented from March 23^rd^, 2020 [41]. This shows that our estimated 95%-CI of NPI dates are in good agreement with the actual dates of NPIs in Germany. Previous study [7] adopting an SIR-type model with weekly reporting modulation and changepoints in transmission rates presents an estimate in transmission rates and change points: *β*_1_ ≈ 0.43→ *β*_2_ ≈ 0.25 → *β*_3_ ≈ 0.15 → *β*_4_ ≈ 0.09 on March 7^th^, March 16^th^ and March 24^th^, respectively. While assuming two changepoints in transmission rate, our inference captured the two major decreases in transmission rate: *β*_1_ ≈ 0.59→ *β*_2_≈ 0.17→ *β*_3_≈ 0.03 on March 6^th^, and March 24^th^, respectively. In comparison, using the two-step methods leads to an overly confident and biased estimation of the model parameters, as the estimated changepoints deviate from the actual NPI dates.

## Discussion

Examples of dynamical systems where model parameters shift at particular points in time abound throughout the sciences. In such cases, inference on both the changepoints themselves and the model parameters is necessary to fully understand the system’s behavior. To date, there has been no formal solution to this important and widespread class of inferential problems. Instead, researchers have had to resort to ad-hoc, two-stage estimation approaches that first use regression methods to detect changepoints, and then treat the changepoints as fixed and known to estimate model parameters on the resulting partitions of the data. While pragmatic, two-stage approaches introduce biases in the estimation of both changepoints and parameters, while also underestimating uncertainty.

In this paper, we developed a formal, simulation-based inference framework for switching dynamical systems with changepoints. Our approach combines models based on mass-action kinetics, which can represent a huge range of complex systems from chemistry, biology, ecology, and physics [22, 42, 43, 44], the LNA as a reliable way to obtain accurate, closed-form likelihood functions for mass-action based models (but see below for limitations), a general approach to computing the gradient of the likelihood function with respect to all parameters (including changepoints) via reverse-mode sensitivity analysis, and Hamiltonian MCMC to efficiently sample the joint posterior distribution of all model parameters (again, including changepoints). The key novelty in our approach lies in adapting and combining these proven components into an integrated framework that supports formal, efficient, and flexible statistical inference for complex systems with parameter discontinuities. Julia-based code repository implementing our approach is also available (see Data Availability).

Using both simulated and real data, we demonstrated the superior performance of our simultaneous estimation framework across a diverse set of examples with contrasting properties and challenges. Specifically, we showed that our approach outperforms two-stage methods in both a best-case scenario for the conventional approach (stochastic gene expression) as well as a more challenging example, with highly oscillatory non-linear dynamics (Lotka-Volterra). In these two examples based on simulated data, the true changepoints and model parameters were known exactly, which allowed us to directly quantify our approach’s superior performance relative to two-stage methods. Additionally, we showed that our method can accurately infer changepoints in a complex epidemiological model applied to limited, noisy, empirical data. In this example, we leveraged the fact that the changepoints are known (the implementation dates of non-pharmaceutical interventions in Germany) to provide an objective point of reference in real-world example. Crucially, this knowledge of NPI implementation dates was used only to check the changepoint estimates produced by our method, and was not used at all in the estimation of model parameters and changepoints.

For likelihood-based inference (including Bayesian methods), obtaining a closed-form likelihood function is intractable for many models. In this work, we used the LNA to overcome this hurdle. Though the LNA is an efficient and effective approximation for the sufficient statistics of stochastic system dynamics [45], this method becomes less accurate when the system has strong nonlinearity plus either when the system size is small, or equivalently when the average number of species is low [46]. In such cases, either higher order correction terms of the LNA [47] or other methods such as moment closure approximations [48], neural-network aided surrogate master equations [49], or invertible neural-network approaches [50] can be considered. A thorough investigation alternative methods of likelihood approximation for more challenging model structures is a clear avenue for further research on our integrated estimation approach.

Another potential advantage of using gradient-based simultaneous optimization that merits further study is its improved accuracy over the approximate Bayesian computation (ABC) method. ABC methods are often used to infer the model parameters of complex dynamical systems when the gradient information is intractable [51, 52, 53], but this flexibility often comes at the cost of being both less accurate and more computationally expensive than likelihood-based approaches. While ABC is not currently widely used for simultaneous inference on switching dynamical systems, we show in the supplementary material (A) that it can indeed be adapted for such purposes. In that predator-prey example, a side-by-side comparison demonstrates our approach clearly outperforms the ABC method in terms of posterior accuracy (see Supplementary Information A for details).

Overall, these findings suggest that our method for simultaneously estimating changepoints and model parameters from switching dynamical systems significantly outperforms conventional two-stage inference methods in terms of estimation accuracy and quantification of uncertainty. The proposed framework shows general applicability to dynamical systems with intrinsic stochasticity and nonlinearity, and we expect that it will become a valuable tool for analyzing a wide range of systems in which both changepoints and model parameters are of primary interest.

## Methods

### Differentiating Through Changepoints and Model Parameters

Consider a system of ODEs with changepoints

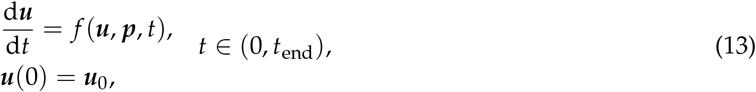

where for fixed *t*, ***u***(*t*) ∈ ℝ^*N*^ and ***p***(*t*) ∈ ℝ^*P*^ are the state and parameter vectors, respectively, and *f* : ℝ^*N*^ *×* ℝ^*P*^ *×* ℝ → ℝ^*N*^. We assume that the parameters ***p***(*t*) are piecewise constant functions, i.e.

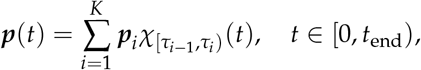

where 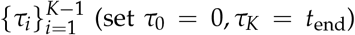 are the changepoints. For simplicity, we denote vector ***θ*** as the concatenation of vectors 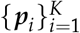 which is of length *PK* and ***τ*** as the concatenation of scalars 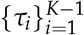 of length *K* − 1. Note that we can also rewrite the right-hand-side of (13) as a function of ***θ*** and ***τ***,

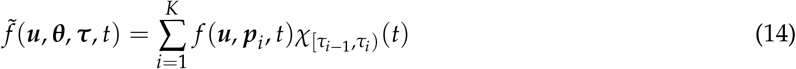

which will be useful for the adjoint equation derivation.

We want to obtain the sensitivity of loss functional *L*(***u, θ, τ***) evaluated over solution ***u***(*t*) = ***u***(*t*, ***θ, τ***)

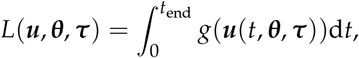

where *g* : ℝ^*N*^ → ℝ. To derive the adjoint sensitivity equations, we introduce the Lagrange multiplier ***λ***(*t*) ∈ ℝ^*N*^ and rewrite

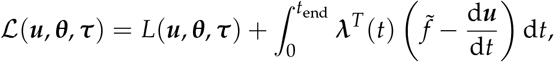

since 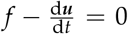 by Equation (13), the sensitivity of parameters to loss functional ℒ is equivalent to the sensitivity of loss functional *L*. Using integration by parts, we have

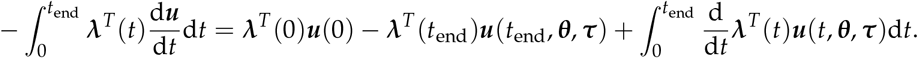

Suppose ***ξ*** is any of the inputs *{****θ, τ, u***_0_*}*,

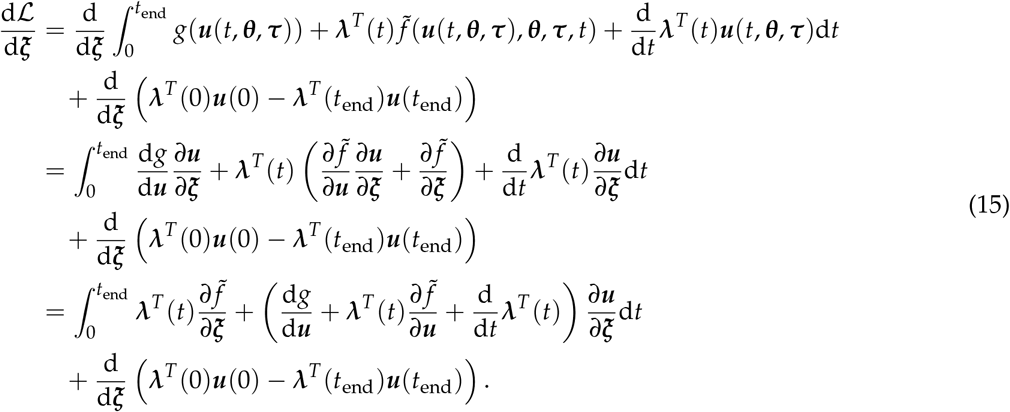

Setting the adjoint equations to satisfy

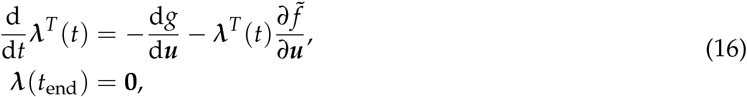

and combining them with Equation (15), leads to the following adjoint equations

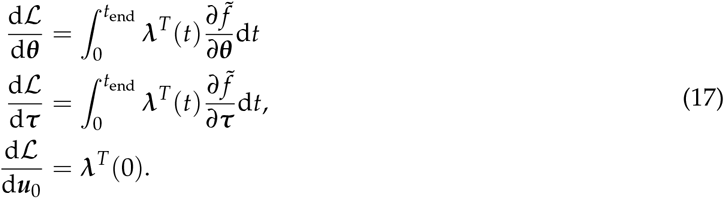

Together with Equation (14), we transform the sensitivity of the system of ODEs with change points (13) which gives

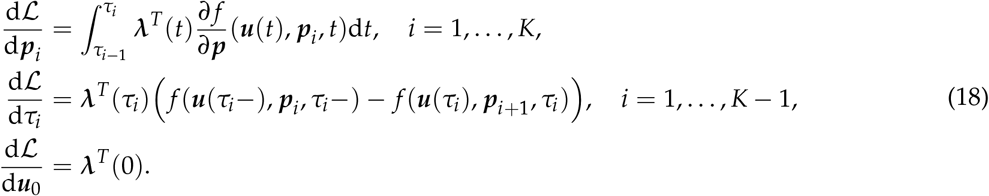

### Constructing the Likelihood Using the Linear Noise Approximation

If an ODE system is based on the law of mass action [25], then the meso-scopic representation (known as stochastic reaction networks) can be used to derive or approximate the likelihood. Such a representation can be mathematically formulated as the MEs, which describe the time evolution of the probability density function of the number of individuals of each species in a well-stirred system. Here we use a moment-based observation model which derives from the LNA. LNA is a widely used approximation method for stochastic reaction kinetics [29].

A general reaction kinetics can be described as follows: given a set of species *X*_*i*_, *i* = 1, …, *N*, we define *R* reactions by the notation

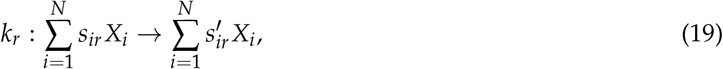

where the stoichiometric coefficients *s*_*ir*_ and 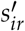 are non-negative integer numbers denoting the numbers of reactant and product molecules, respectively. The quantity *k*_*r*_ in Equation (19) is called the reaction rate constant of the *r*th reaction. Classically, the dynamics of a reaction system as in Equation (19) is modelled by the law of mass action. The law of mass action states that the rate of a reaction is proportional to the product of the concentrations of reactant molecules, which lead to the following rate equations as:

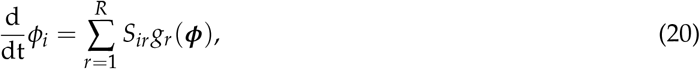

where *ϕ*_*i*_ is the concentration of species *X*_*i*_,

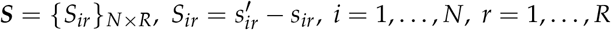

is the stoichiometric matrix, and

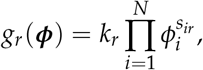

is the rate of the *r*th reaction.

However, the law of mass action is only valid when the number of molecules is large. When the number of molecules is small, System (19) can instead be modelled by a continuous-time Markov jump process to study the probability of the system being in a particular state at a given time. The dynamics of such a system can be described by the MEs [30]:

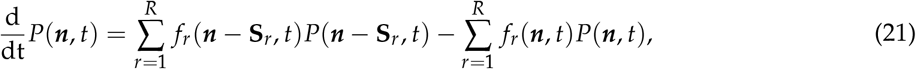

where *P*(***n***, *t*) is the probability of the system being in state ***n*** at time *t*,

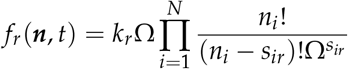

is the propensity function of reaction *r* at state ***n***, and **S**_*r*_ is the stoichiometric vector of reaction *r*, Ω is the system size (or volume of the system).

The MEs are written directly from the rate constants and stoichiometries of all the elementary reactions, but neither analytical nor numerical solutions are in general available. Fortunately, the MEs can often be simplified in an LNA. Linear noise approximation is an expansion of the CME taking the inverse system size 1/Ω as the perturbed variable, which is originally developed by [54, 29]. The idea is to separate concentrations into a deterministic part, given by the solution of the deterministic rate equations, and a part describing the fluctuations about the deterministic mean

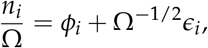

where *ϕ*_*i*_ is the solution of the deterministic rate equations (20), and *ϵ*_*i*_ represents fluctuations about the deterministic mean. Define **σ** = (*ϵ*_*ij*_)_*N×N*_ to be the covariance matrix of the fluctuations, the LNA is given by:

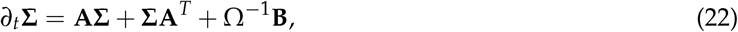

where **A** = **A**(***ϕ***), **B** = **B**(***ϕ***) both depend on the solution ***ϕ*** of the rate Equation (20), which are defined by

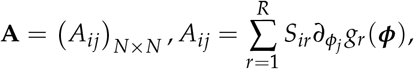

as the Jacobian matrix of the deterministic rate equations, and 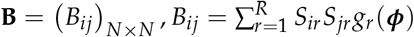.

In this formulation, the LNA allows for analytical solutions that are locally valid close to macroscopic trajectories (solution of the rate equations) of the system. We refer to the review paper [30] for more details on the CME and LNA.

Once we have system (22), we can construct the likelihood function directly by assuming it follows a normal distribution. Specifically, if for species *i*, the observed data is denoted as 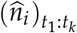 at time points {*t*_1_, …, *t*_*k*_}, the mean and variance of the species *i* are given by solving *ϕ*_*i*_ and *ϵ*_*ii*_ using coupled Equations (20) and (22) simultaneously.

In practice, it is also common to modify the variance of the LNA to account for the practical measurement errors, for example, the variance of the LNA can be rescaled by *σ*^2^ for the noise intensity parameter *σ*. Then we define the following likelihood as the observation model:

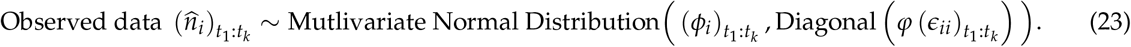

Here *φ* represents a function that maps the variance of the LNA to the variance of the measurement errors.

### Bayesian Inference with Hamiltonian Monte Carlo

Our Bayesian inference uses sampling based on a Hamiltonian Monte Carlo (HMC) algorithm. HMC is a Markov Chain Monte Carlo (MCMC) algorithm that uses Hamiltonian dynamics to propose new states for sampling from a probability distribution. It involves simulating a physical system with a Hamiltonian function, where the position and momentum of a fictitious particle are used to represent the current state of the system. HMC requires the gradient of the log probability density function to provide the force needed to update the variables of interest through the potential energy landscape of the distribution, which allows for more efficient exploration of the parameter space compared to traditional MCMC methods.

The No-U-Turn Sampler (NUTS) is an extension of the Hamiltonian Monte Carlo algorithm. NUTS eliminates the need for users to specify parameters such as the number of steps *L* required for each iteration. This is because NUTS uses a recursive algorithm that automatically decides when to stop the iteration. Additionally, NUTS adapts the step size *ε* during the sampling process using a dual averaging algorithm, which enables more efficient exploration of the target distribution. However, NUTS, like other types of Hamiltonian Monte Carlo sampling methods, requires gradient information from the log probability density function (pdf) of the target distribution. This is because the newly proposed sample is generated by integrating the Hamiltonian dynamics:

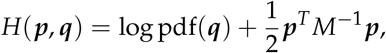

where ***q*** is the position variable (the variable of our target distribution) and ***p*** is the momentum variable, which is introduced artificially to make the Hamiltonian dynamics to work properly, *M* is the mass matrix, and *H* is the Hamiltonian function. To obtain the newly proposed sample, we first initialize the position variable ***q*** and the momentum variable ***p***, and then we integrate the Hamiltonian dynamics using the leapfrog algorithm [55, 56]

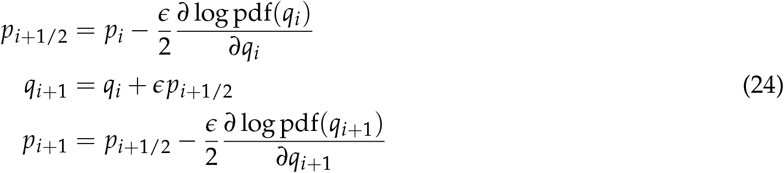

where *ϵ* is the step size, and 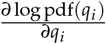 are the gradients of the log probability density function evaluated at *q*_*i*_.

Thanks to the sensitivity analysis on both change points and model parameters, the gradient of the log probability density function is available using automatic differentiation techniques. With this, we can use NUTS to sample from the posterior distribution of the model parameters.

### Software for Numerical Experiments

In our numerical experiments, we use the *Julia* programming language [57]. We use *LinearNoiseApproximations*.*jl* [58] for LNA analysis. We adapt the package *SciMLSensitivity*.*jl* and *DifferentialEquations*.*jl* [59] to perform the sensitivity analysis, and use the package *Turing*.*jl* [60] to perform the Bayesian inference.

## Data Availability

The original the number of reported COVID-19 cases (infected and deceased) in Germany can be obtained from [39]. The synthetic data and the code for this paper is available at https://github.com/xiaomingfu2013/SwitchDyn-Changepoints-Parameters-Identification.

## Supplementary Information

### A Lotka-Volterra (Predator-Prey) Model

In this example, we consider is the Lotka-Volterra model of predator and prey interactions. The corresponding ODE system is given by

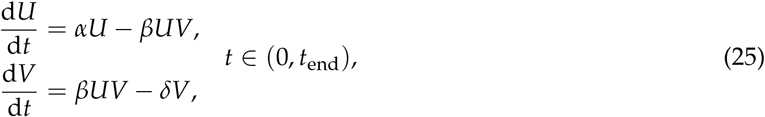

with initial conditions *U*(0) = *U*_0_, *V*(0) = *V*_0_, where *α* represents the growth rate of the prey population in the absence of predator, *β* is the interaction strength between the predator and prey populations, and *δ* is the mortality rate of the predator population. The corresponding reaction network is given by

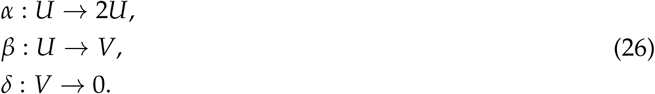

The growth rate of prey population *α* (similar to the interaction rate *β*) are assumed to be piecewise constant functions, that is

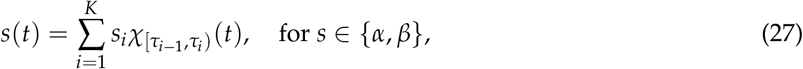

where 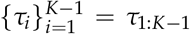 are the changepoints and 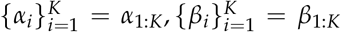 are the corresponding piecewise rates. We also estimate the mortality rate *δ*, but assume its value is constant over time.

Following the linear noise approximation (LNA) method in the main text Equation (22), the expanded (co)variance solution from the LNA model of the system (25) is given by

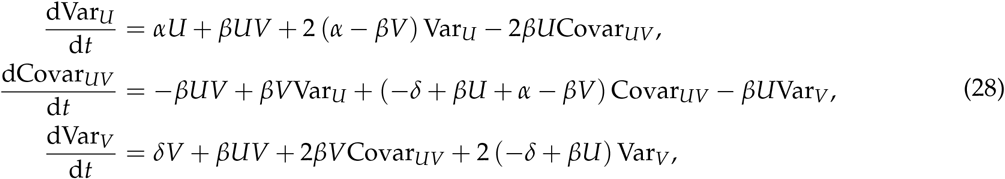

where we set the system size Ω = 1 in the main text Equation (22).

In our implementation, we set the number of changepoints to one and the initial conditions for prey and predator population to *U*_0_ = 120, *V*_0_ = 140 respectively. A total number of 300 sets of model parameters are randomly drawn from the sample distributions in Table 2. The synthetic data is generated from the model (25) and (28) for each parameter set. To mimic the measurement errors, the generated synthetic data are further perturbed by noise following Normal 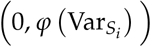 where *S*_*i*_ and 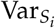 are the population and its variance at time *t*_*i*_ and 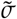 is the noise intensity for *S* ∈ *{U, V}*, respectively. We set the number of time snapshots to *N* = 500 taken from time interval to (0, 50) equidistantly. We set the noise intensity 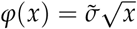 with 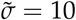.

**Table 2:** Parameter description of model (25)

| Parameter | Description | Sample Distributions |
| --- | --- | --- |
| $\alpha_1$ | First growth rate of the prey population | $\text{LogNormal}(\log(0.8), 0.1)$ |
| $\alpha_2$ | Second growth rate of the prey population | $\text{LogNormal}(\log(0.6), 0.1)$ |
| $\beta_1$ | First interaction rate between the predator and prey | $\text{LogNormal}(\log(5 \times 10^{-3}), 0.1)$ |
| $\beta_2$ | Second interaction rate between the predator and prey | $\text{LogNormal}(\log(3 \times 10^{-3}), 0.1)$ |
| $\delta$ | Mortality rate of the predator population | Fixed to 0.4 |
| $\tau$ | Change point | $\text{Uniform}(22, 28)$ |

**Table 3:** Comparison of the posterior distributions of the model parameters between our method and the ABC method.

| Symbols | True | Mean (Ours) | Mean (ABC) | Std. (Ours) | Std. (ABC) | Median (Ours) | Median (ABC) |
| --- | --- | --- | --- | --- | --- | --- | --- |
| $\alpha_1$ | 0.82 | 0.83 | 0.82 | $4.01 \times 10^{-3}$ | 0.02 | 0.83 | 0.82 |
| $\alpha_2$ | 0.62 | 0.62 | 0.63 | $7.16 \times 10^{-3}$ | 0.04 | 0.62 | 0.63 |
| $\beta_1$ | $4.71 \times 10^{-3}$ | $4.72 \times 10^{-3}$ | $4.76 \times 10^{-3}$ | $2.03 \times 10^{-5}$ | $1.22 \times 10^{-4}$ | $4.72 \times 10^{-3}$ | $4.76 \times 10^{-3}$ |
| $\beta_2$ | $4.00 \times 10^{-3}$ | $3.99 \times 10^{-3}$ | $4.00 \times 10^{-3}$ | $5.45 \times 10^{-5}$ | $2.38 \times 10^{-4}$ | $4.00 \times 10^{-3}$ | $4.00 \times 10^{-3}$ |
| $\tau$ | 24.93 | 24.68 | 28.20 | 0.22 | 4.88 | 24.68 | 26.97 |

For the inference of the model parameters and simulation-based calibration, we assume no prior knowledge on the changepoint (i.e. we set the prior distribution of the changepoint *τ*∼ Uniform(0, 50)). We use the observation model given by

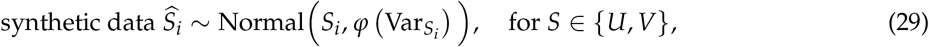

where *S*_*i*_ (and 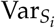) are the population (and its variance) at time *t*_*i*_ from Equation (25) and (28). We use the No-U-Turn Sampler (NUTS) to sample from the posterior distribution of the model parameters with accepted rate of 0.65, and total sample length of 500 with 250 adaptation (warm-up) iterations.

To check the ability of our Bayesian model to recover the correct target posterior, also referred as computational faithfulness, we use simulation-based calibration (SBC) [31]. SBC is a diagnostic method which considers the performance of a sampling algorithm over the entire Bayesian joint distribution, regardless of the structure of the particular model. It leverages the fact that most Bayesian models are generative by construction. The brief procedure is described as follows: First, we generate random draws from the prior distribution 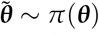 and the corresponding synthetic data 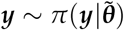 from the model. We can then build inferences for each simulated observation 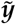 and compare the recovered posterior distribution to the sampled parameter 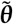. If a Bayesian sampling method generates samples from the exact posterior, the following equality should hold regardless of the particular form of the posterior

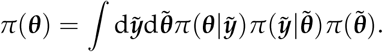

Thus, any deviation from this equality suggests that an error has been introduced by the sampling method. The error could be caused by imprecise computation of the posterior or an erroneous implementation of the model itself.

In practice, we draw 300 sets of random parameters from the prior distributions (see Supplementary Information A). Each sampled parameter configuration is then used to run the model forward and generate synthetic predator-prey trajectories. The posterior distribution is then computed using Bayesian inference on each synthetic dataset. All the rank statistics are within the 99% confidence intervals, which is consistent with the expectation of uniformity.

Figure 4 **a** shows the SBC based on comparing histograms of rank statistics to the discrete uniform distribution. The rank statistics are computed by sorting the posterior samples and assigning a rank to each sample. The rank statistics are then normalized to the range [0, 1]. The histograms of the rank statistics are expected to be uniform if the posterior samples are drawn from the exact posterior distribution [31], thus proving the computational faithfulness of the sampling method. We can see that the gray shade in marks the 99% confidence intervals of the uniform histograms. All the rank statistics are within these confidence intervals, which is consistent with the expectation of uniformity.

**Figure 4:**
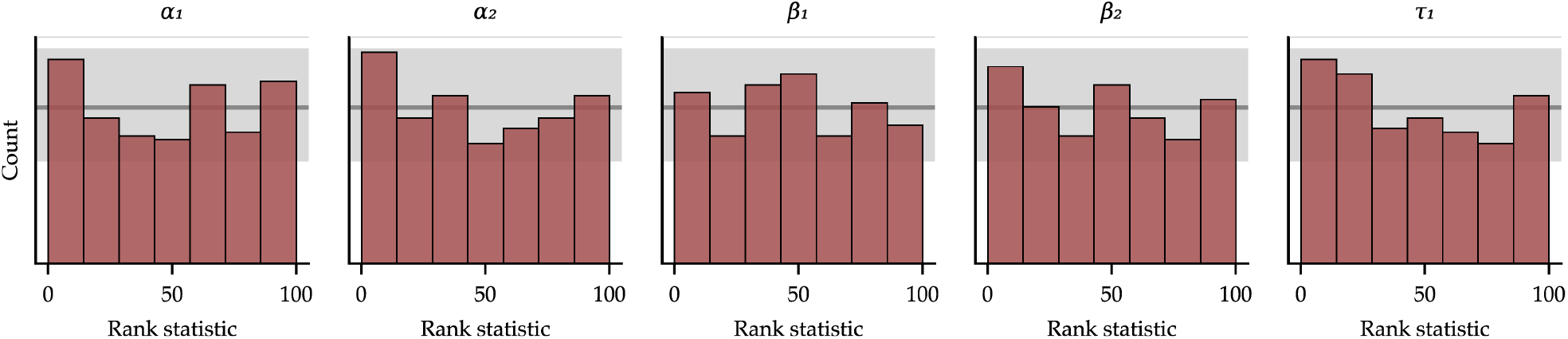
Uncertainty calibration on results with the Lotka-Volterra model. Simulation-based calibration (SBC) computed from 300 simulated predator-prey scenarios. The uniformity of the rank statistics indicates that there are no systematic biases in the approximate posteriors and implies trustworthy approximation.

#### Comparison with Approximate Bayesian Computation (ABC)

Here we showcase the comparison of using our inference method and the approximate Bayesian computation (ABC) method. ABC methods are often used to infer the model parameters of complex dynamic systems when the likelihood function is intractable [51, 52]. However, the ABC method is known to be less efficient at obtaining an accurate posterior distribution [52]. We compare the inference results on the same synthetic dataset used in the predator-prey model in Figure 1. For the ABC method, we used the ABC sequential Monte Carlo (ABC-SMC) method [51] with the same prior distributions in Table 2. We set the final number of particles to 10^3^, the *α* quantile for the next generation to 0.2, and the maximum function evaluations to 10^6^. In Figure 5, we can see that the posterior distributions of the model parameters using our method are more concentrated and closer to the true values than the ABC method. The summary statistics of the posterior distributions in Table A also show that both the mean and median of the two methods are close to the true values, while the standard deviations using our method are much smaller than the ABC method.

**Figure 5:**
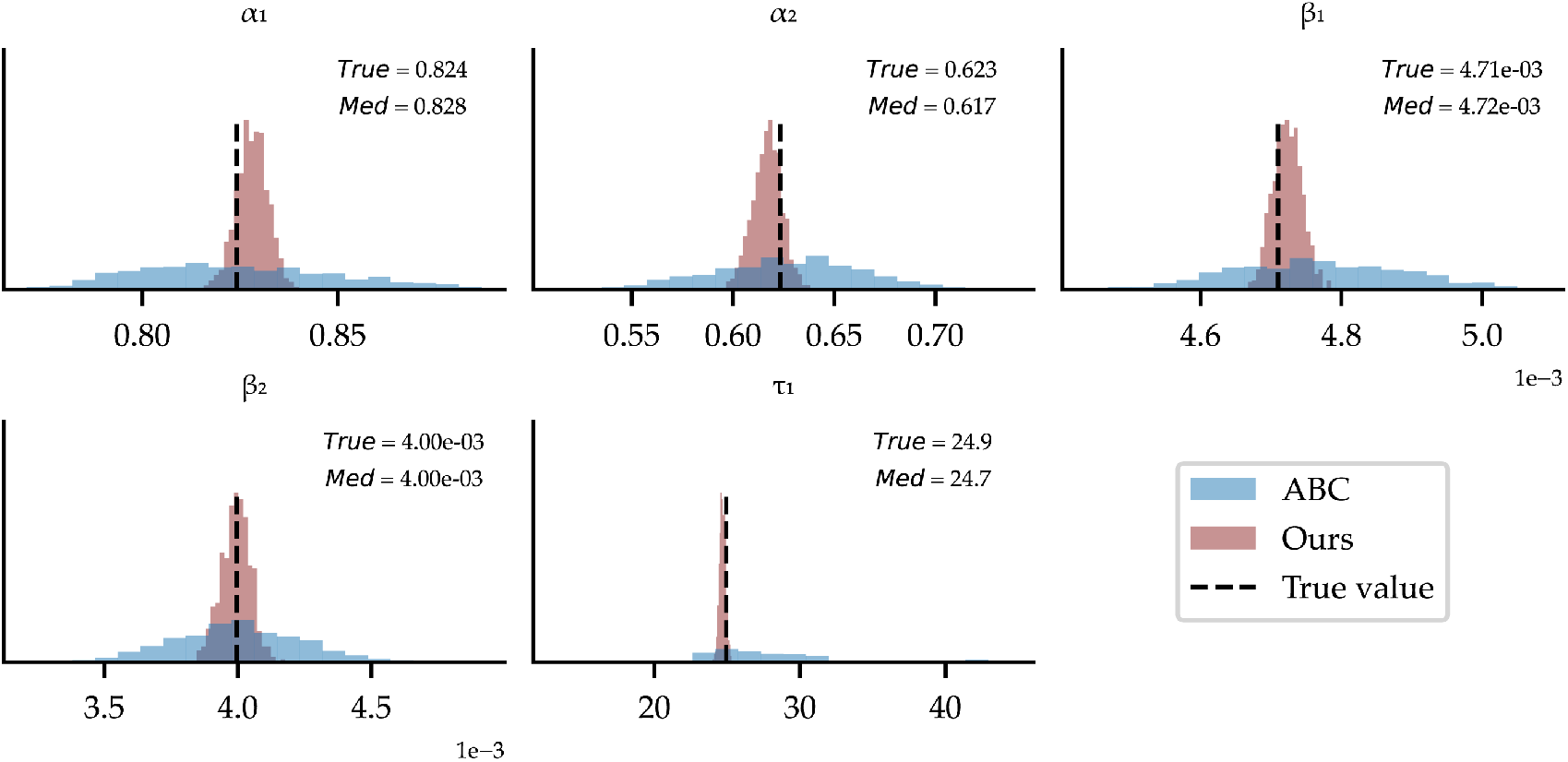
Comparison of the inferred posterior distributions of the model parameters using our method and the ABC method. The true values are shown as the vertical dashed lines.

### B Stochastic Gene Expression Model

In this example, we consider a simple stochastic gene expression model with changepoints. The corresponding reaction network is given by

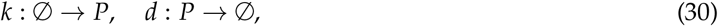

where *P* represents the protein, *k* represents the initiation rate, and *d* represents the degradation rate. The corresponding MEs are given by

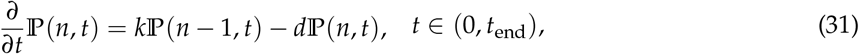

where ℙ(*n, t*) is the probability that the system is in state *n* at time *t*. The production rate *k* is assumed to change at the changepoints *τ*, that is

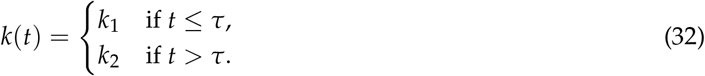

It is well-known that during a cell cycle the underlying dynamics of stochastic gene expression can undergo a phase transition, thus affecting the effective initiation rate *k*. However, it is also observed that such transitions are asynchronous in a population, that is, the transitions happen at different times with different initiation rates for different cells. In this example, we introduce the heterogeneity of the cell dynamics by assuming that, for each cell, the two initiation rates before and after the changepoint *τ* are random variables and the changepoint *τ* also follows a random distribution. The distributions of *k*_1_, *k*_2_ and *τ* are given by

For this simple example with a changepoint, one can derive that the solution of the MEs (31) follows a Poisson distribution, that is,

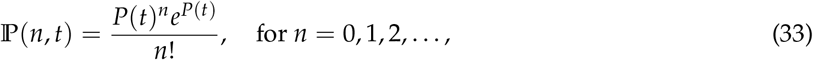

where *P*(*t*) is the solution of the corresponding ODE system

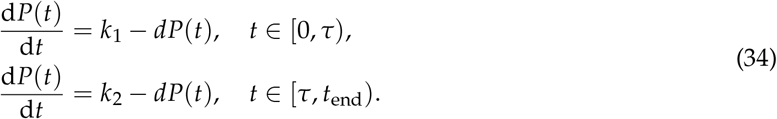

For generating the synthetic data, we use the stochastic simulation algorithm (SSA) [36] to simulate 300 sets of random trajectories from (30) with parameters drawn from random distributions in Table 4. We fix the initial condition *P*(0) = 5. Then for each parameter set *j* = 1, …, 300, our inference task is to accurately estimate the initiation rate 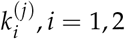 and the changepoint *τ*^(*j*)^ using Bayesian inference with the negative log-likelihood function (33),

**Table 4:** Parameter description of model (31)

| Parameter | Description | Parameter Distributions |
| --- | --- | --- |
| $k_1$ | First initiation rate | LogNormal(log(30), 0.1) |
| $k_2$ | Second initiation rate | LogNormal(log(15), 0.1) |
| $\tau$ | Changepoint | LogNormal(log(15), 0.1) |
| $d$ | degradation rate of the predator population | Fixed to 1 |

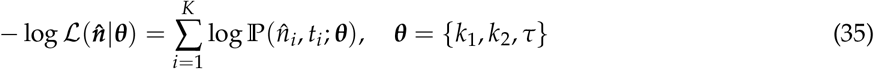

where 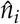 is the observed protein number at time *t*_*i*_ and *K* is the total number of observations.

### C Compartmental (SEIRD) Model

In this example, we consider a SEIRD model including the susceptible (S), exposed (E), infected (I), recovered (R) and deceased (D) compartments. The corresponding ODE system is given by

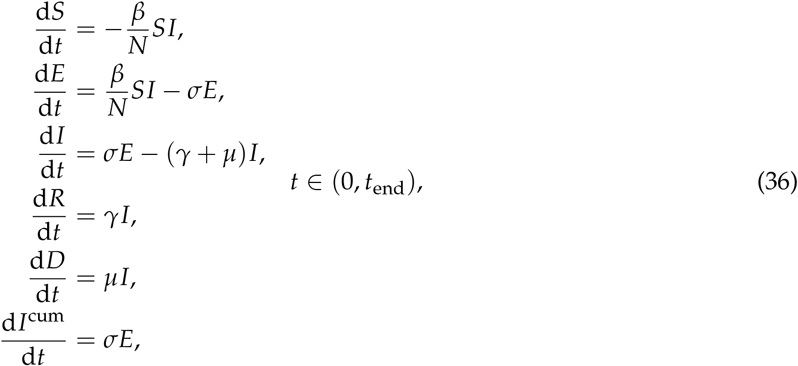

where *β* is the transmission rate, *γ* is the recovery rate and *N* = *S* + *E* + *I* + *R* is the total number of the population excluding the deceased group. The parameter *σ* is the rate at which exposed individuals become infectious, and *μ* is the rate at which infected individuals die. Here *I*^cum^ is an auxiliary compartment to record the cumulative infected number for comparison with observed cumulative infected data.

The initial conditions *I*(0), *D*(0) are given by the observed data, while the initial condition *E*(0) is determined by *I*(0)(1 + *σ*/*γ*) where *σ*/*γ* is the ratio of the incubation period to the average infectious period, and *R*(0) = 0. The initial condition *S*(0) is determined by *N*− (*I*(0) + *E*(0) + *R*(0) + *D*(0)), where *N* is the total population of Germany. Finally, we normalize the population size to population per million for the parameter inference. The meaning of the model parameters and their priors are detailed in Table 5. The prior distributions and parameters are chosen and adapted based on the study [7, 61].

**Table 5:** Parameter description of model (36).

| Parameter | Description | Prior Distributions |
| --- | --- | --- |
| $\beta_1$ | First transmission rate | LogNormal(log(0.5), 0.2) |
| $\beta_2$ | Second transmission rate | LogNormal(log(0.25), 0.2) |
| $\beta_3$ | Third transmission rate | LogNormal(log(0.05), 0.2) |
| $\mu_1$ | First mortality rate | LogNormal(log($1 \times 10^{-3}$), 0.1) |
| $\mu_2$ | Second mortality rate | LogNormal(log($4 \times 10^{-3}$), 0.1) |
| $\sigma$ | Rate at which exposed cases become infectious | LogNormal(log(0.25), 0.1) |
| $\gamma$ | Recover rate | Fixed to 1/14 |
| $\tau_{\beta_1}$ | First changepoint for the transmission rate | Uniform(10, 35) |
| $\tau_{\beta_2}$ | Second changepoint for the transmission rate | Uniform(35, 60) |
| $\tau_{\mu_1}$ | First changepoint for the mortality rate | Uniform(10, 60) |

The corresponding reaction network of Equation (36) can be written as

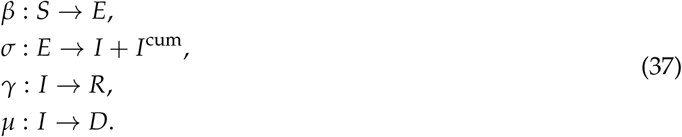

Similarly, using Methods section Constructing the Likelihood Using the Linear Noise Approximation for the derivation LNA of the SEIRD model, we obtain the coupled SEIRD model with the time-dependent (co)variance solution of each group (for simplicity we omit the recover group *R*)

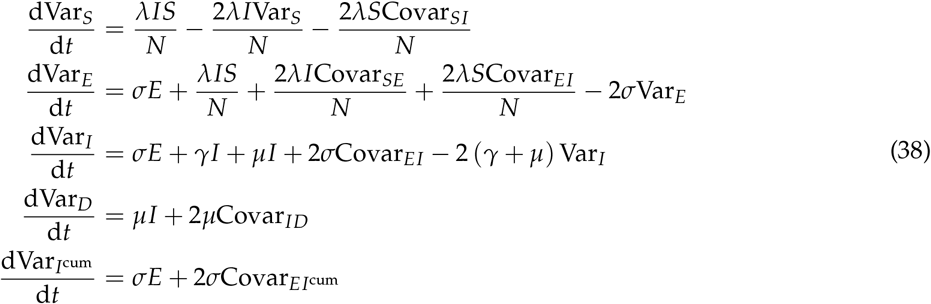

and the coupled covariance equations

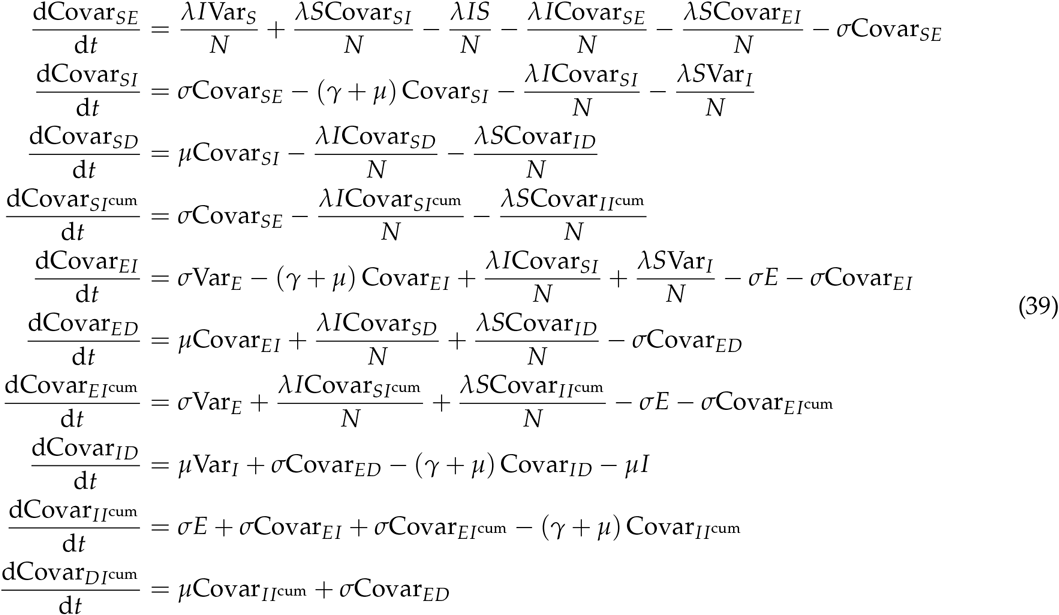

Since the observed cumulative data has only small fluctuations, we scale the variance of LNA by *σ*^2^ = 10^−2^. Thus, if the observation data is given by a time-series on 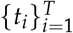 denoted as 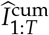 and 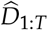, the likelihood function reads

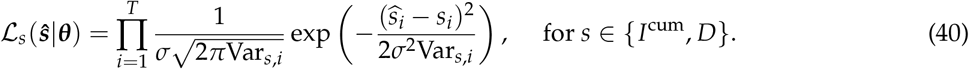

Maximizing the likelihood function with respect to the parameters ***θ*** is equivalent to minimizing the negative log-likelihood function

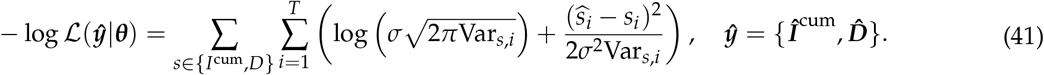

For the inference of the parameters, we use the NUTS to sample from the posterior distribution of the model parameters with accepted rate of 0.65, and total sample length of 1000 with 500 adaptation iterations.

#### Verification of the Inferential Accuracy on Synthetic Data

Likewise, to check computational faithfulness, we use the SBC and compare the estimated median of posterior samples to the ground truth to verify the inferential accuracy. In practice, we use the synthetic data generated from 300 sets of parameters drawn from the prior distributions (Table 5) using model (36). To avoid the corner case where the interventions are too close to each other, we set the prior of change points to be

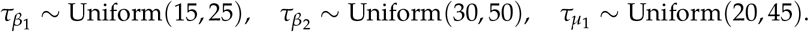

The corresponding synthetic data is perturbed by Gaussian noise according to Equation (40). The inference is done using LNA according to Equations (38)-(40). The SBC results are shown in Figure 6 **a** and the inferential accuracy in Figure 6 **b**. We can see that the uniformity of the rank statistics indicates that the inference is faithful to the model (Figure 6 **a**), and the estimated parameters are close to the ground truth, with the error bars marking the median absolute deviation of the posterior samples (Figure 6 **b**).

**Figure 6:**
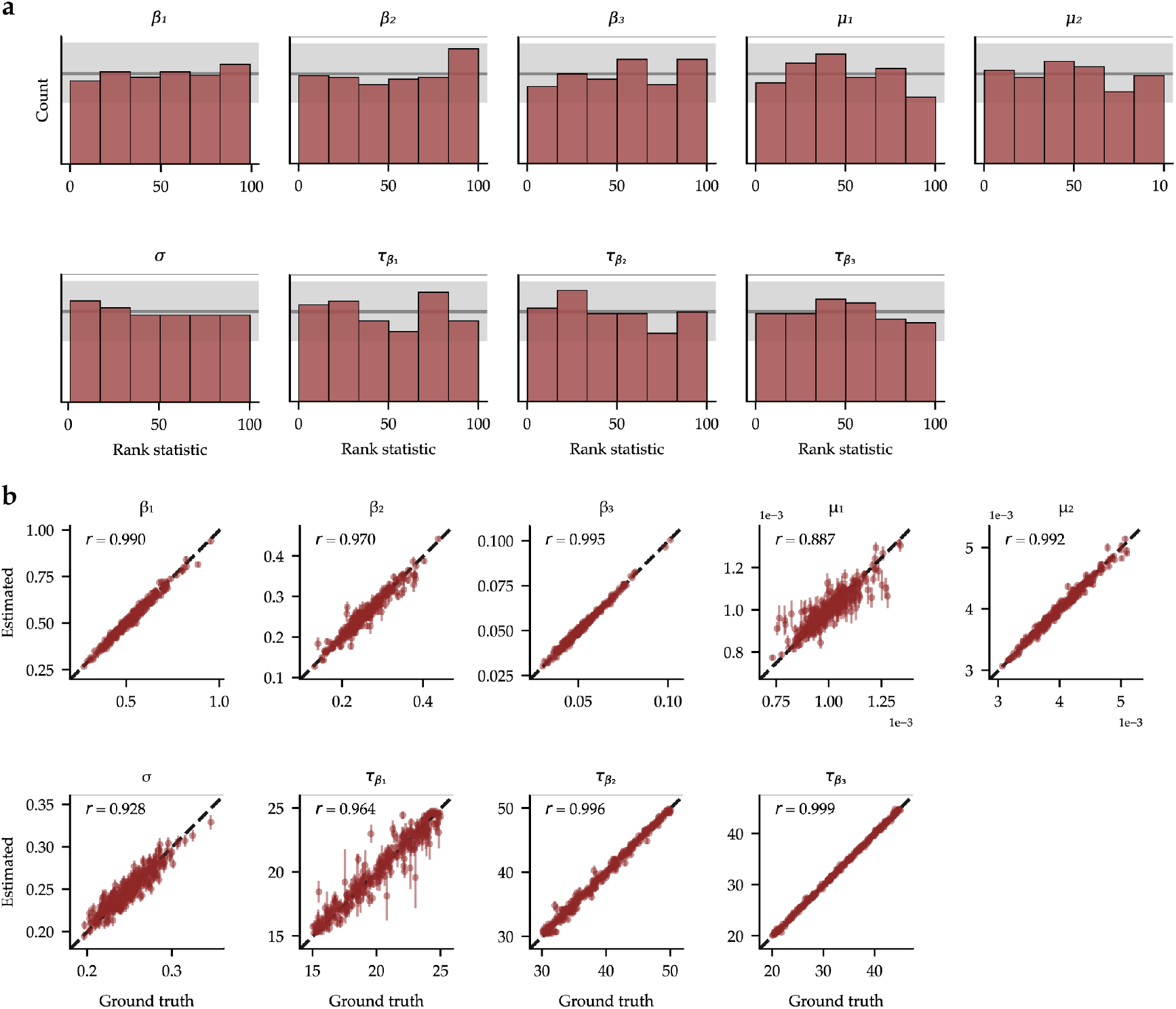
**a**. Simulation-based calibration (SBC) results of the SEIRD model from 300 sets of synthetic data. The uniformity of the rank statistics indicates that the inference is faithful to the model. **b**. Inferential accuracy of the SEIRD model by reusing the synthetic data. The error bars mark the median absolute deviation of the posterior samples.

#### Inference Results for Two Sets of Change Points in Mortality Rate

In the main text, we presumed two sets of change points for transmission rate *β* and one set of change point for mortality rate *μ*. Here, we add an additional change point to the mortality rate *μ* to see if the inference results are sensitive to the number of change points *τ*_*μ*_. The prior distributions of the model parameters are the same as in Table 5, except that we add additional priors for the third mortality rate *μ*_3_ and the second change point 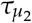, that is

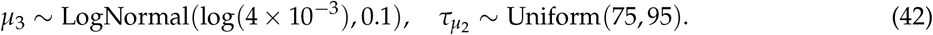

Compared to the results in Figure 3, we can see from Figure 7 that the posterior distributions of the third mortality rate *μ*_3_ and the second change point 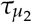 have very limited changes compared to the priors, suggesting that one change point 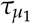 for the mortality rate *μ* might be sufficient. This is also supported by the inference results in Table 6, where the inferred parameters are close to the results in Table 1.

**Table 6:** Parameter inference results of the Germany data with two sets of changepoints for transmission rate and mortality rate.

| Description | Parameter | Mean | Median | MAP | 95%-CI |
| --- | --- | --- | --- | --- | --- |
| Transmission rate $S \rightarrow E$ (1/days) | $\beta_1$ | 0.59 | 0.59 | 0.59 | [0.55, 0.62] |
| | $\beta_2$ | 0.16 | 0.16 | 0.16 | [0.15, 0.18] |
| | $\beta_3$ | 0.03 | 0.03 | 0.03 | [0.03, 0.04] |
| Mortality rate $I \rightarrow D$ ($1 \times 10^{-3}$/days) | $\mu_1$ | 1.02 | 1.02 | 1.03 | [0.92, 1.10] |
| | $\mu_2$ | 3.89 | 3.89 | 3.89 | [3.79, 3.99] |
| | $\mu_3$ | 3.85 | 3.81 | 3.71 | [3.26, 4.55] |
| Incubation rate $E \rightarrow I$ (1/days) | $\sigma$ | 0.29 | 0.29 | 0.29 | [0.27, 0.31] |
| Changepoint for the transmission rate | $\tau_{\beta_1}$ | Mar-21<br>(Day 20.35) | Mar-21<br>(Day 20.34) | Mar-21<br>(Day 20.26) | [Mar-20, Mar-22]<br>[Day 19.49 - 21.19] |
| | $\tau_{\beta_2}$ | Apr-09<br>(Day 38.66) | Apr-09<br>(Day 38.62) | Apr-09<br>(Day 38.62) | [Apr-07, Apr-10]<br>[Day 37.14 - 40.49] |
| Changepoint for the mortality rate | $\tau_{\mu_1}$ | Mar-28<br>(Day 27.22) | Mar-28<br>(Day 27.34) | Mar-28<br>(Day 27.48) | [Mar-27, Mar-29]<br>[Day 26.46 - 27.80] |
| | $\tau_{\mu_2}$ | May-25<br>(Day 85.08) | May-25<br>(Day 85.08) | May-27<br>(Day 87.38) | [May-16, Jun-04]<br>[Day 75.52 - 94.65] |

**Figure 7:**
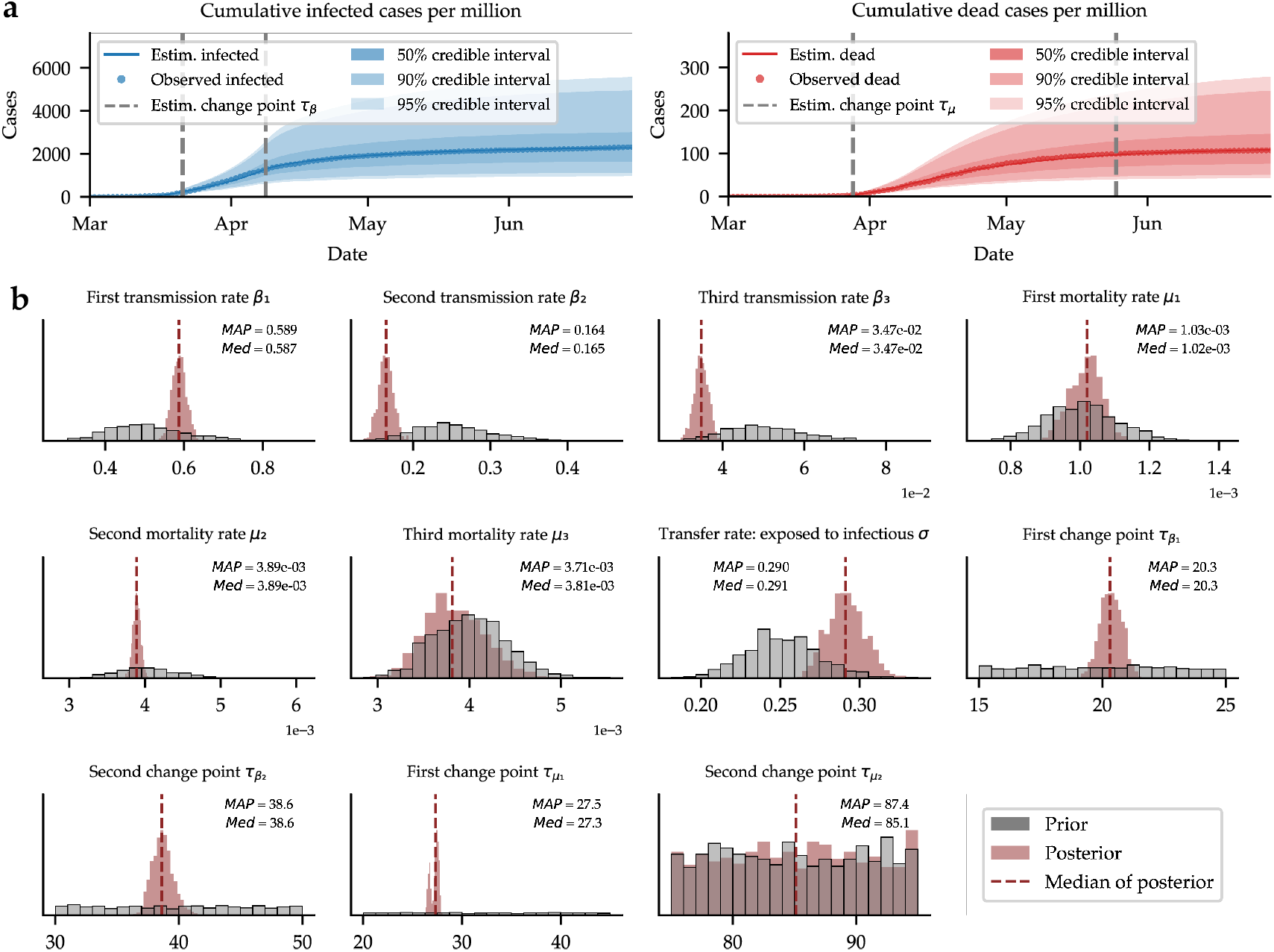
**a**. Posterior predictions of new cases obtained by inferring model parameters from epidemiological data available for reported infected, and dead. **b**. Marginal posteriors of all model parameters inferred from data alongside median and MAP summary statistics. Gray histograms depict prior distributions for comparison with the posteriors. Vertical dashed lines indicate posterior medians. For comparison, the posterior distributions of the mortality rate *μ*_3_ and the second changepoint 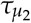 show very limited changes compared to the priors.

### D Comparison with Two-stage Inference Methods

In comparison to our simultaneous inference method, we use a conventional two-stage parameter inference method, namely detecting the changepoint first and then estimating model parameters in a second step as the changepoint is fixed and known. We use four distinct R packages that implement different changepoint detection algorithms: *segmented* [62], *strucchange* [18], *Rbeast* [19], and *mcp* [63, 64, 20]. While both *segmented* and *strucchange* rely on (generalized) linear models, they differ in their approach, with *segmented* requiring fitted lines to connect estimated changepoints through regression models, and *strucchange* using generalized fluctuation tests and F tests for detection. In contrast, both *Rbeast* and *mcp* employ Bayesian regression for changepoint detection. *Rbeast* decomposes the trend and seasonality in the time-series for reliable changepoint detection, while *mcp* uses JAGS [65] (a variation of Gibbs sampler) for changepoint detection, allowing regression models to differ between segments.

#### Lotka-Volterra (Predator-Prey) Model

In the predator-prey system, we use the same 300 sets of synthetic data in the main text section Case Study: Lotka-Volterra (Predator-Prey) Model. Instead of setting the changepoint *τ* as a variable to be inferred, we use the four R packages to detect the changepoint first. Among the four detection methods, *mcp* demands the longest computation time, while its parameter estimation has the least relative error (e.g. |*τ*_inferred_ − *τ*_true_ |/*τ*_true_) compared the assumed ground truth as demonstrated in our case studies (see Figure 8).

**Figure 8:**
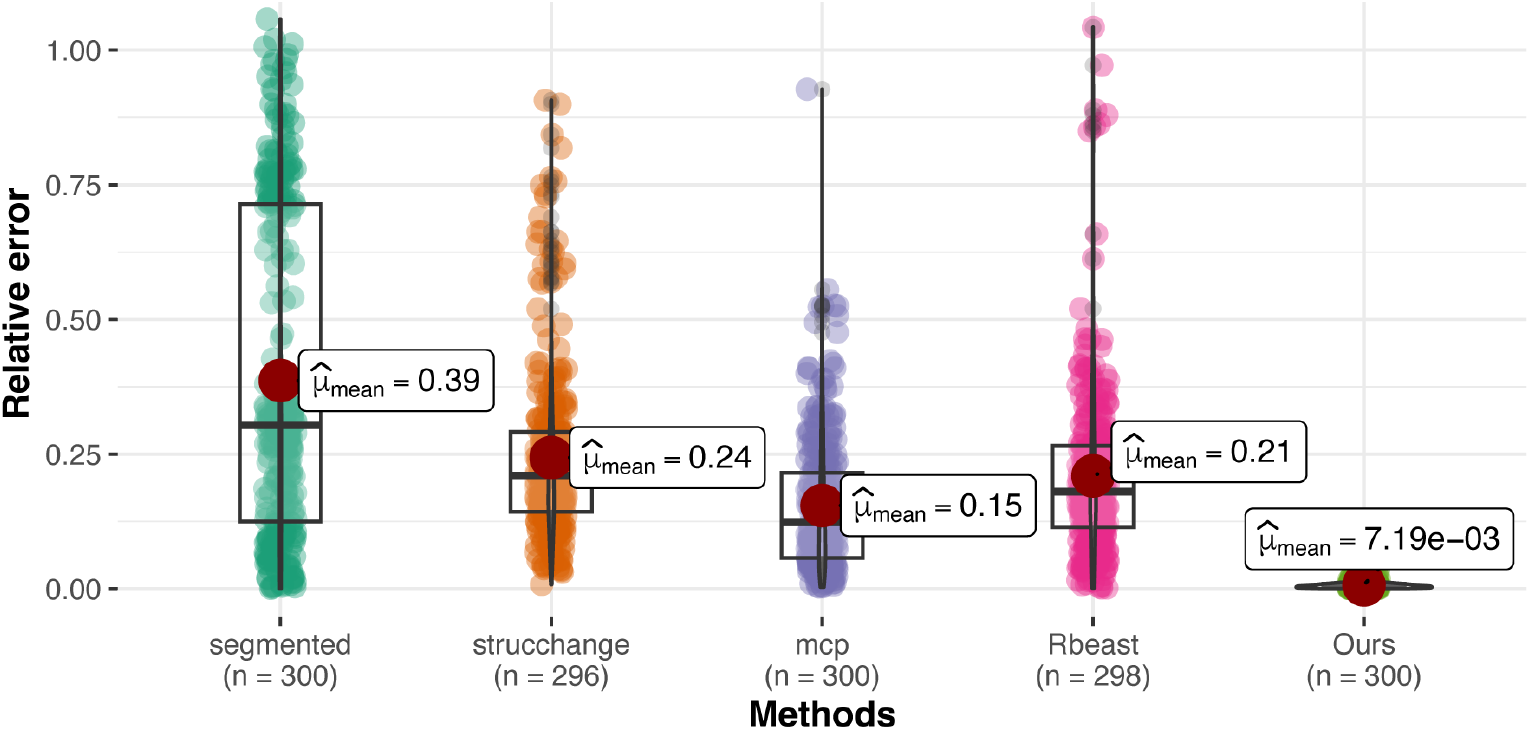
We compare the inferred changepoints from the four R packages with the ground truth. The relative error of the inferred changepoints are shown in the violin plot. For Methods section *strucchange* and *Rbeast* the valid sample numbers are 296, and 298 respectively, as the algorithms fail to detect changepoints in the remaining cases.

Then we choose the results of the most accurate change point detection method *mcp* and use the inferred change points to estimate rest of the model parameters with the same model-based inference procedure as ours over the 300 sets. The inferred parameter distributions are shown in Figure 1. We can see that the overall inferred parameters from the two-stage methods are much less accurate than our integrated inference. This is because once the changepoint detection is not accurately inferred, the posteriors of rest of the parameters are not reliable. See Figure 9 for a typical example. While the inferred posteriors using the two-stage method exhibit strong concentration, they diverge significantly from the ground truth.

**Figure 9:**
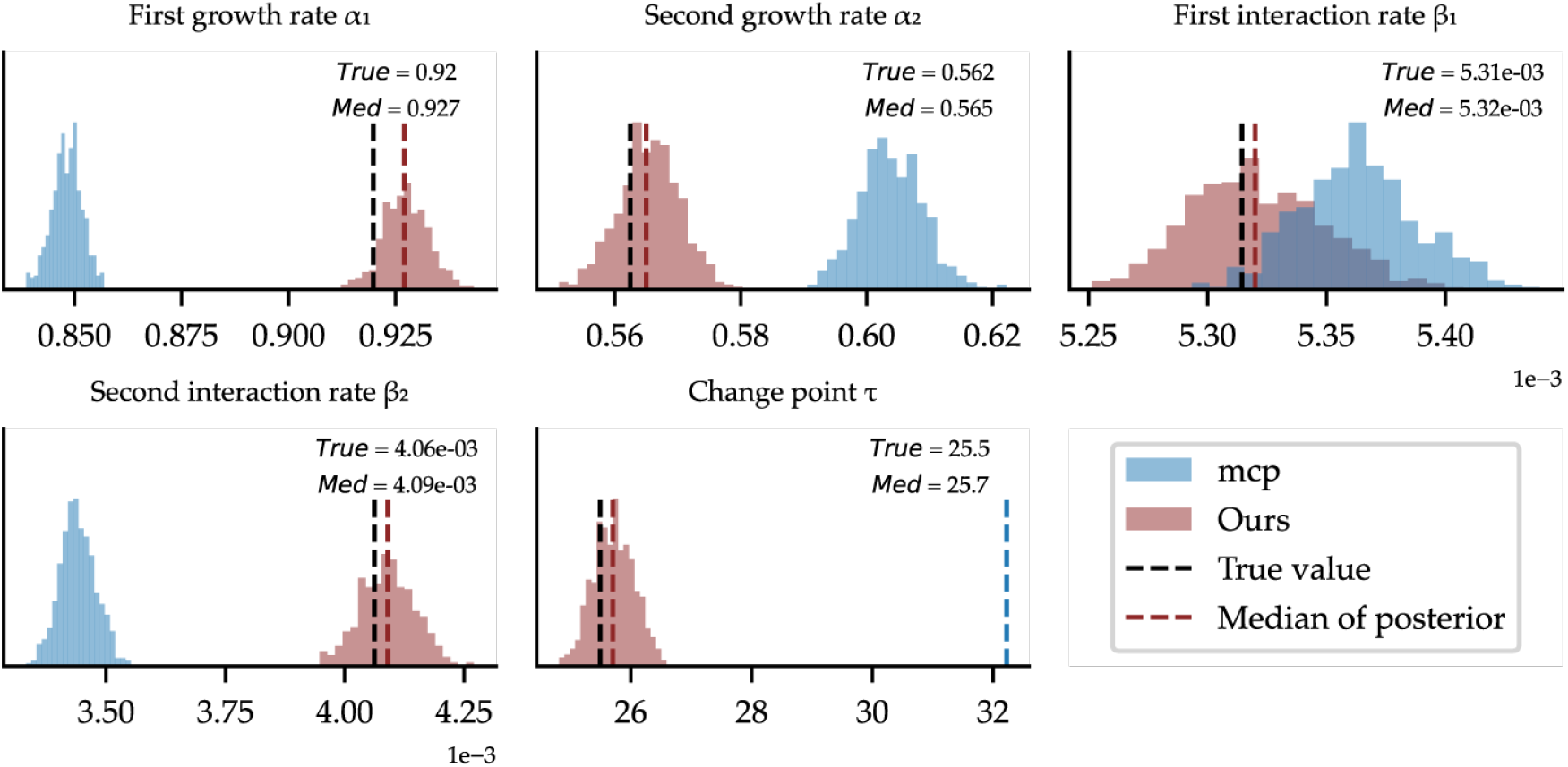
One representative posterior plot out from 300 synthetic cases: our method versus the two-stage method based on *mcp*. While the inferred posteriors using the two-stage method exhibit strong concentration, they diverge significantly from the ground truth.

The details of the inferred parameters are shown in Table 7.

**Table 7:** Parameter inference results of the Lotka-Volterra in Figure 9.

| Parameter | Description | True | Median (Ours) | Median (mcp) |
| --- | --- | --- | --- | --- |
| $\alpha_1$ | First growth rate of the prey population | 0.92 | 0.93 | 0.85 |
| $\alpha_2$ | Second growth rate of the prey population | 0.56 | 0.57 | 0.60 |
| $\beta_1$ | First interaction rate between the predator and prey | $5.31 \times 10^{-3}$ | $5.32 \times 10^{-3}$ | $5.36 \times 10^{-3}$ |
| $\beta_2$ | Second interaction rate between the predator and prey | $4.06 \times 10^{-3}$ | $4.09 \times 10^{-3}$ | $3.44 \times 10^{-3}$ |
| $\delta$ | Mortality rate of the predator population | 0.4 (fixed) | - | - |
| $\tau$ | Changepoint | 25.49 | 25.71 | 32.22 |

**Table 8:** Comparison of the inferred parameters (median statistics) and the negative log-likelihood using the two-stage method and our integrated inference method. The negative log-likelihood is calculated using the median of the inferred parameters.

| Parameter | Description | segmented | strucchange | mcp | Rbeast | Ours |
| --- | --- | --- | --- | --- | --- | --- |
| $\beta_1$ | First transmission rate | 1.08 | 0.38 | 0.38 | 1.08 | 0.59 |
| $\beta_2$ | Second transmission rate | 0.13 | 0.03 | 0.03 | 0.12 | 0.16 |
| $\beta_3$ | Third transmission rate | 8.39e-03 | 0.02 | 0.02 | 8.03e-03 | 0.03 |
| $\mu_1$ | First mortality rate | 8.03e-04 | 2.14e-03 | 2.17e-03 | 2.97e-03 | 1.02e-03 |
| $\mu_2$ | Second mortality rate | 3.40e-03 | 6.69e-03 | 6.74e-03 | 4.09e-03 | 3.89e-03 |
| $\sigma$ | Transfer rate: exposed to infectious | 0.16 | 0.53 | 0.52 | 0.15 | 0.29 |
| $\gamma$ | Recover rate (fixed) | 0.07 | 0.07 | 0.07 | 0.07 | 0.07 |
| $\tau_{\beta_1}$ | First changepoint for transmission rate | 17.30 | 31.00 | 30.86 | 18.00 | 20.53 |
| $\tau_{\beta_2}$ | Second changepoint for transmission rate | 51.59 | 52.00 | 51.15 | 52.00 | 38.67 |
| $\tau_{\mu_1}$ | First changepoint for mortality rate | 13.29 | 50.00 | 50.55 | 51.00 | 27.46 |
| $\mathcal{L}$ | Negative log-likelihood | 1795.07 | 1509.29 | 1520.34 | 1802.50 | **892.11** |

#### Stochastic Gene Expression Model

Similar to the predator-prey system, we use the same 300 sets of synthetic data in the stochastic gene expression model in section Case Study: Stochastic Gene Expression Model. We use the four R packages to detect the changepoint first. Among the four detection methods, *strucchange* performs the best in terms of the relative error of the inferred changepoints (see Figure 10).

**Figure 10:**
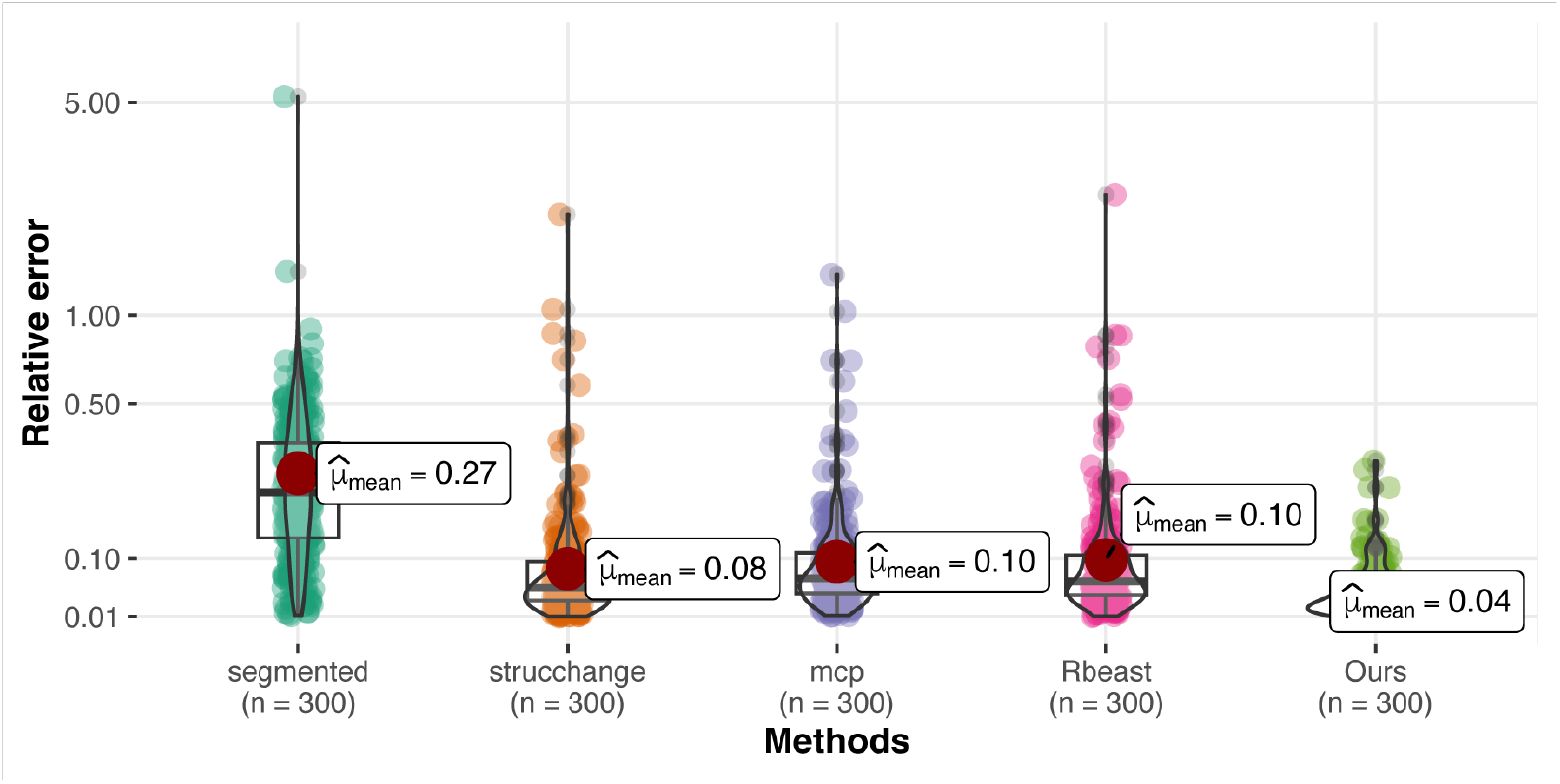
We compare the inferred changepoints from the four R packages with the ground truth. The relative error of the inferred changepoints are shown in the violin plot.

Under the conditions that are more favorable to two-step method, the mean relative error for the inferred changepoints *strucchange* is 0.08, while ours is still two times smaller (0.04). Then we choose the results of the most accurate changepoint detection method *strucchange* and use the inferred change points to estimate rest of the model parameters with the same model-based inference procedure as ours over the 300 sets. The full results are presented in Figure 2.

#### Compartmental Models in Epidemiology

In a comparison study, we consider the previous two-stage parameter inference method, that is to identify the changepoint using the four changepoint detection methods and then to estimate model parameters using the same inference settings as our inference method. The inference results using four different methods are shown in Figures 11, 12, 13 and 14 respectively. We can see that the inferred parameters from the two-stage method are less accurate than our integrated inference method. This can be also confirmed by the negative likelihood value using the median of the inferred parameters (Table D).

**Figure 11:**
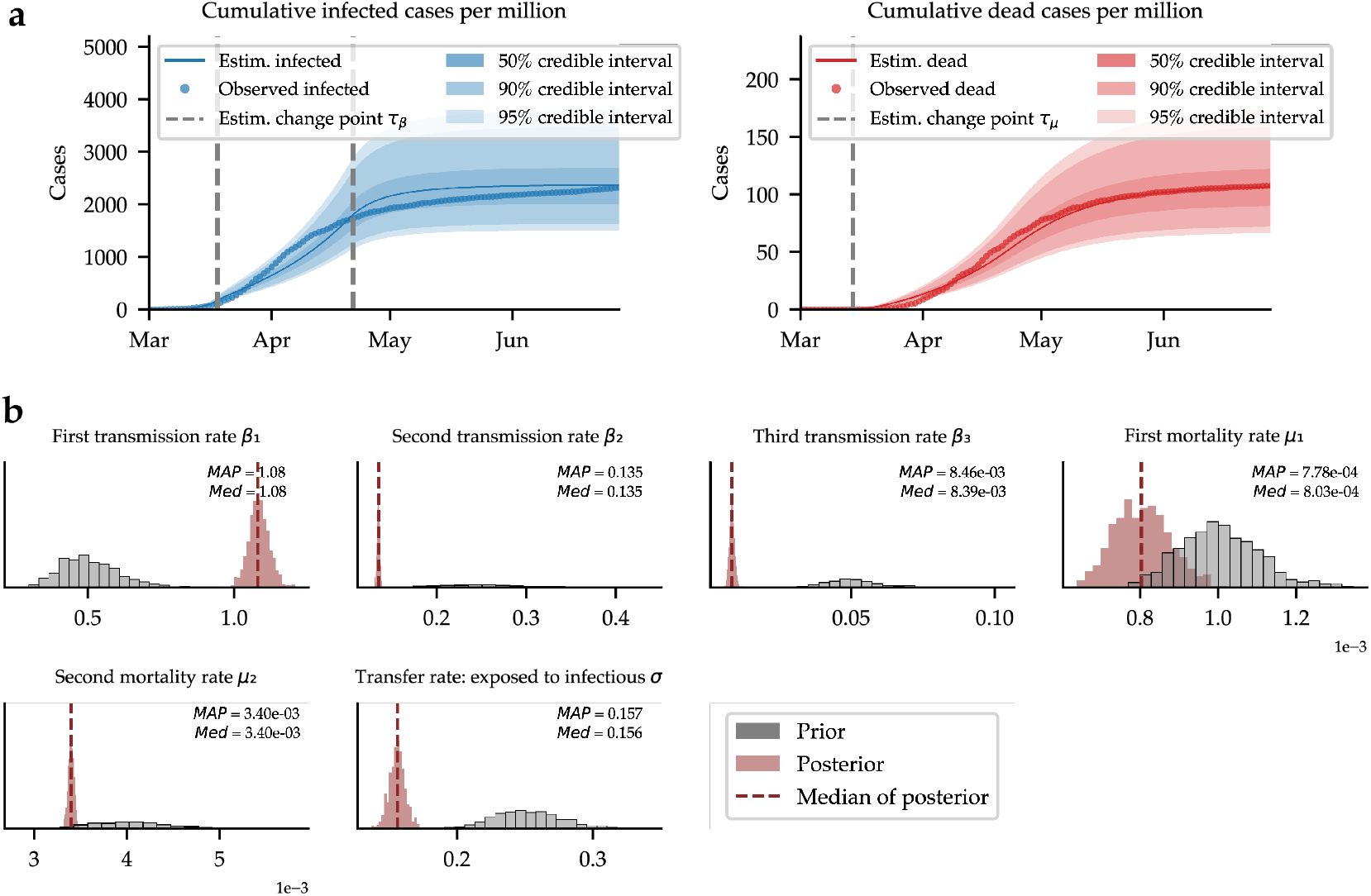
Inference results using the two-stage method based on *segmented*. **a**. Posterior predictions of new cases obtained by inferring model parameters from epidemiological data available for reported infected, and dead. **b**. Marginal posteriors of all model parameters inferred from data alongside median and MAP summary statistics. Gray histograms depict prior distributions for comparison with the posteriors.

**Figure 12:**
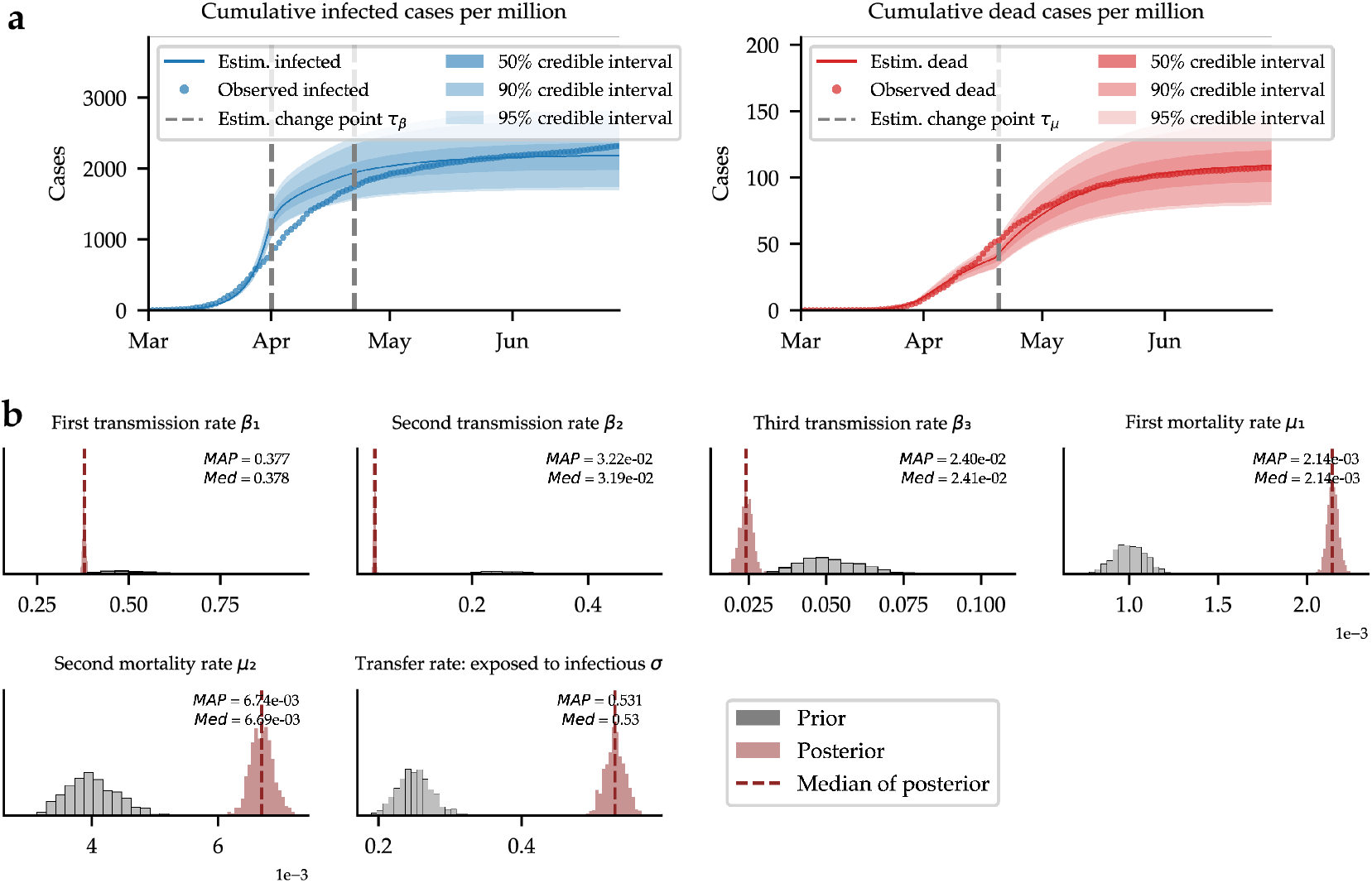
Inference results using the two-stage method based on *strucchange*. **a**. Posterior predictions of new cases obtained by inferring model parameters from epidemiological data available for reported infected, and dead. **b**. Marginal posteriors of all model parameters inferred from data alongside median and MAP summary statistics. Gray histograms depict prior distributions for comparison with the posteriors.

**Figure 13:**
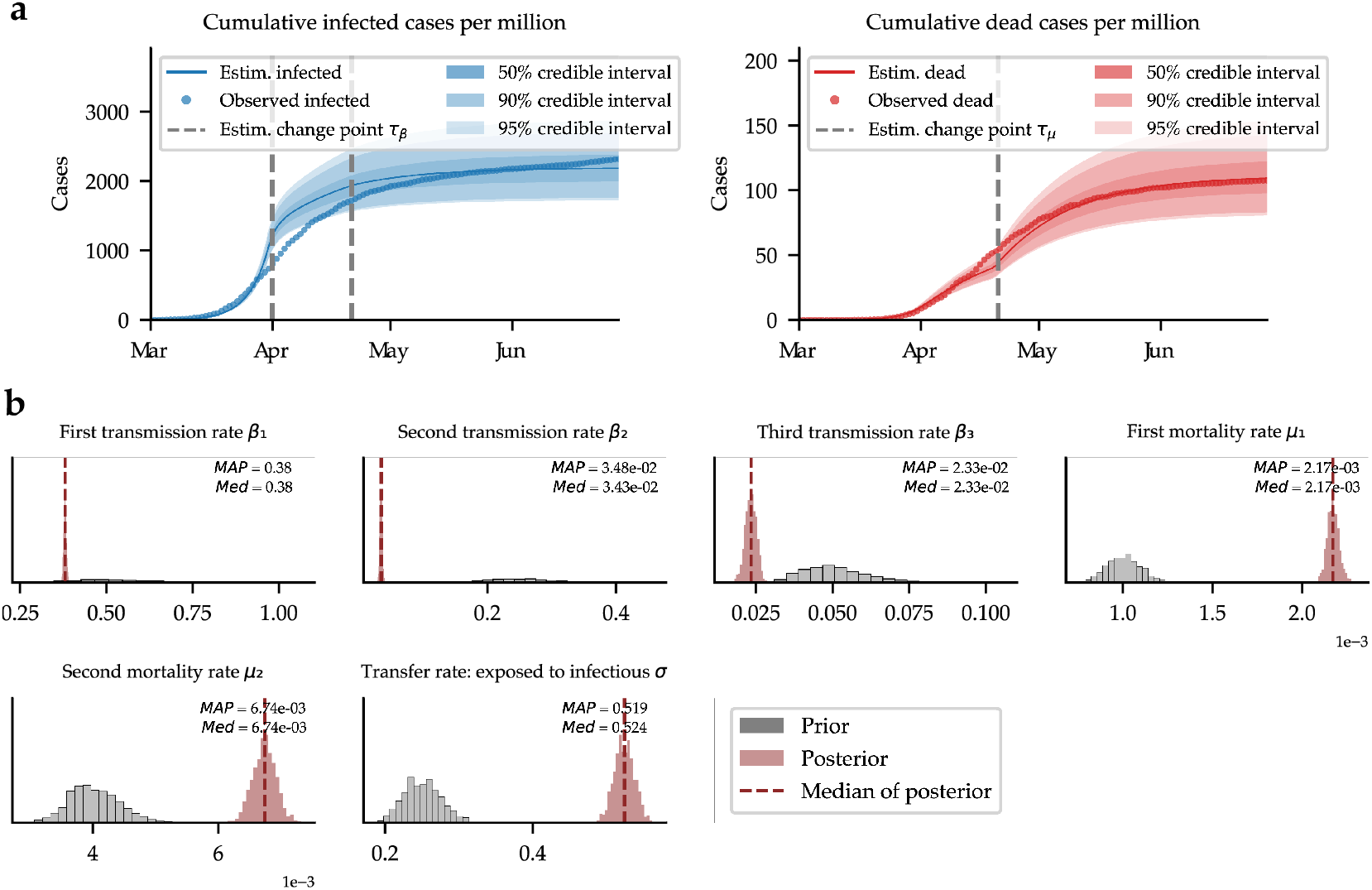
Inference results using the two-stage method based on *mcp*. **a**. Posterior predictions of new cases obtained by inferring model parameters from epidemiological data available for reported infected, and dead. **b**. Marginal posteriors of all model parameters inferred from data alongside median and MAP summary statistics. Gray histograms depict prior distributions for comparison with the posteriors.

**Figure 14:**
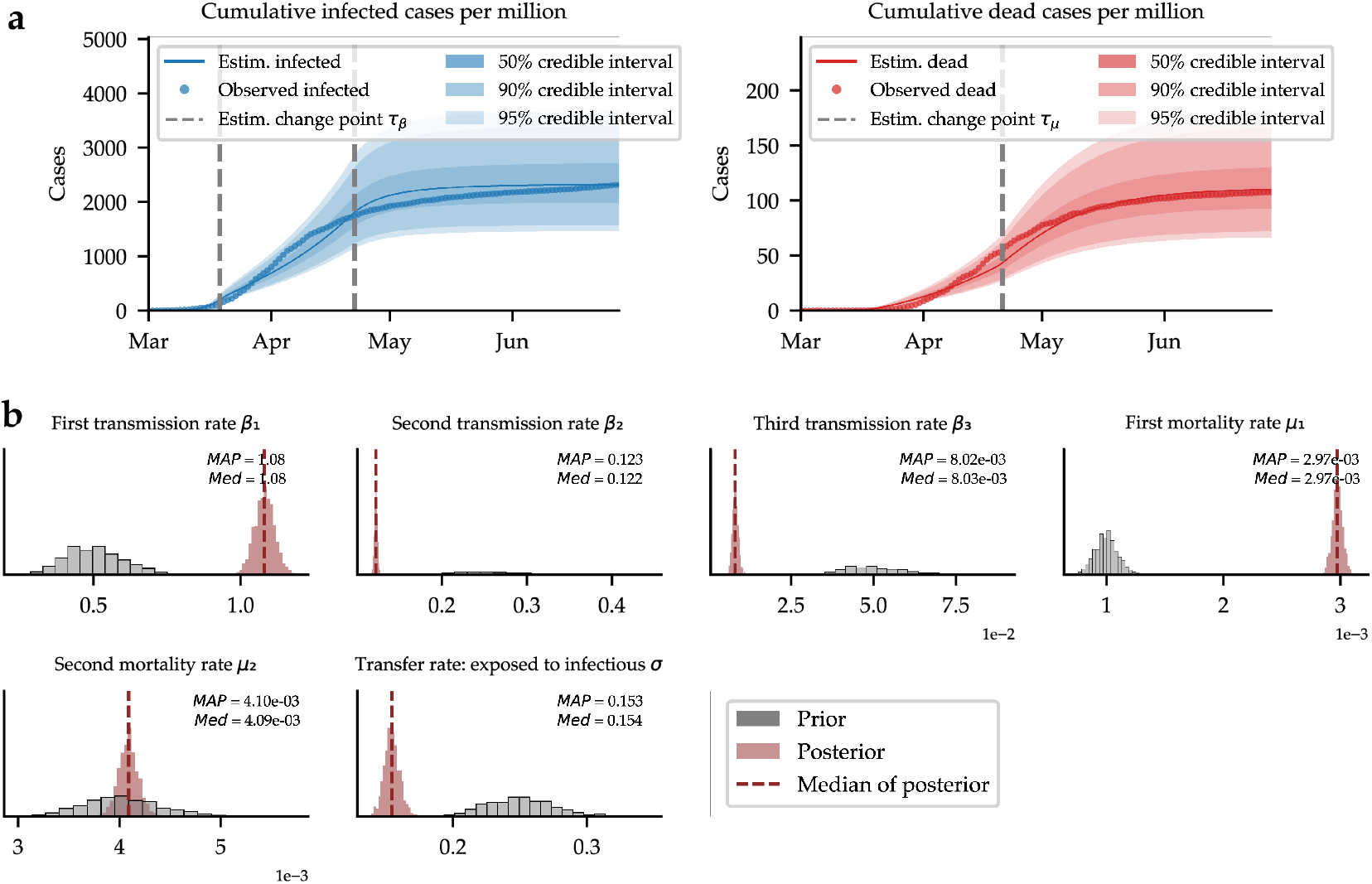
Inference results using the two-stage method based on *Rbeast*. **a**. Posterior predictions of new cases obtained by inferring model parameters from epidemiological data available for reported infected, and dead. **b**. Marginal posteriors of all model parameters inferred from data alongside median and MAP summary statistics. Gray histograms depict prior distributions for comparison with the posteriors.

